# A Translational Platform for Brain-Computer Interfaces and Adaptive Neuromodulation: Technical Characterization, Long-Term Validation, and Implementation of the CorTec Brain Interchange–BCI2000 Ecosystem

**DOI:** 10.64898/2026.08.27.747359

**Authors:** Frederik Lampert, Matthew R Baker, Filip Mivalt, William Engelhardt, Nicholas Luczak, Alexis Gkogkidis, Martin Schüttler, Amir H Ayyoubi, Behrang Fazli Besheli, Max A van den Boom, Jordan Bilderbeek, Douglas J Kellar, Inyong Kim, Vaclav Kremen, Nathan P Staff, Gerwin Schalk, Nuri F Ince, Peter Brunner, Gregory A Worrell, Kai J Miller

## Abstract

**Objective:** Adaptive neuromodulation systems and implantable brain-computer interfaces (BCIs) are promising therapies for neurological and psychiatric disorders. However, their broader translation into research and clinical practice remains limited by technological complexity, restricted access to implantable research platforms, and the lack of standardized, reproducible experimental workflows. We therefore aimed to develop and validate an open, general-purpose translational ecosystem that enables rapid development, evaluation, and dissemination of novel neuromodulation and implantable BCI paradigms.

**Approach:** The CorTec Brain Interchange (BIC) implantable neural sensing and stimulation device was integrated with the open-source BCI2000 platform to create a modular, extensible neuromodulation ecosystem. We established a standardized battery of quantitative assessments to characterize implantable neuromodulation systems to comprehensively evaluate the CorTec BIC device through benchtop characterization, long-term preclinical in vitro and in vivo validation, and a human proof-of-concept demonstration.

**Results:** Benchtop and saline testing provided a comprehensive technical ex vivo characterization of the BIC device, independently validating previously reported performance while extending its characterization through quantification of the recording noise floor, stimulation and acquisition latencies and impedance measurement accuracy. Long-term in vivo validation in five canines, with the longest implantation exceeding three years, demonstrated stable chronic recordings while capturing progressive channel deterioration and its underlying mechanical causes. The ecosystem enabled active functional decoding more than two years after implantation, implementation of closed-loop stimulation using arbitrary spectral biomarkers, detection and modulation of epilepsy-associated biomarkers, and brain stimulation evoked potential recordings. In addition, we translated an established one-dimensional BCI cursor control paradigm to the BIC benchtop evaluation kit and demonstrated its feasibility in a human participant. Finally, we openly provide standardized surgical, © Year Copyright holder imaging, and analysis pipelines together with datasets and software to facilitate reproducible neuromodulation research.

**Significance:** We present a versatile, open-source translational ecosystem that supports a wide range of neuromodulation and implantable BCI applications with minimal modification. This battery of quantitative assessments can be applied generally as a blueprint for systematic characterization of implantable neuromodulation systems. By combining comprehensive hardware characterization with standardized software tools and experimental workflows, this work provides both an essential reference for researchers adopting the Brain Interchange platform. The ecosystem lowers technical barriers to implantable neurotechnology research, promotes reproducibility, and provides a foundation for accelerating the development and clinical translation of next-generation adaptive neuromodulation and implantable BCI therapies for patients with neurological and psychiatric disorders.

## 1. Introduction

Neurological and psychiatric disorders affect millions worldwide and are leading causes of ill health and disability worldwide [1], [2], [3]. While pharmacologic treatments exist for many of these disorders, they often lack efficacy in some patients or have serious side-effects [4], [5].

Neuromodulation is a promising therapeutic option for patients with medication-resistant disorders, offering targeted, modulable and reversible therapies [4], [5], [6], [7], [8], [9]. Continuous open-loop neuromodulation is an established therapy for several movement and epilepsy disorders [7], [9].

More recently, adaptive neuromodulation has demonstrated the potential to improve therapeutic outcomes by tailoring stimulation to ongoing neural activity, enabling more targeted and responsive therapy. This approach has shown promise in movement disorders [10], [11], [12], stroke rehabilitation [13], [14], psychiatric disorders [15] [16], [17], [18], and spinal cord injury [19], [20]. Adaptation of stimulation parameters can occur across multiple temporal scales, ranging from fast closed-loop control on the order of tens of milliseconds, through responsive stimulation operating over seconds [10], [20], [21], to longer-term adaptation that accounts for changes occurring over hours such as circadian cycles [5], [22], [23] or fluctuations in medication levels [24].

Understanding the temporal dynamics of the nervous system and the ability to modulate neural activity across multiple time scales remain major challenges in neuromodulation therapies [25], [26]. Although considerable progress has been made in this area [22], [25], [26], [27], current research is largely confined either to prolonged in-hospital monitoring [23], [28] or to studies employing implantable neural sensing and stimulation devices [27], [29], [30].While hospital-based recordings provide access to high-bandwidth neural data, they cannot capture the diversity of behaviors and environments encountered during daily life. Conversely, current implantable systems enable chronic recordings in naturalistic settings but remain constrained by limited sampling rates, channel counts, onboard computation, and processing flexibility. As a result, sophisticated adaptive stimulation algorithms and investigation of neural dynamics at high temporal and spatial resolution remain difficult to implement outside controlled clinical environments [31], [32].

In parallel, the field of brain-computer interfaces (BCIs) has experienced rapid technological progress and substantial financial investment in recent years [33], [34], leading to several landmark demonstrations of chronically implanted BCI systems [20], [35], [36], [37], [38]. Despite these advances, translation into broadly applicable clinical therapies has remained limited [39], [40] [41]. Many of the most advanced implantable BCI systems have been developed through successful collaborations between academia and industry, resulting in remarkable technological advances.

However, these systems typically remain accessible to only a small number of research centers, limiting broader community participation, independent validation, and widespread dissemination of methods and knowledge [39].

We believe that broader adoption of open, standardized research ecosystems can lower the technical barriers to entry in implantable neurotechnology research and accelerate clinical translation through more reproducible and accessible research. This is further reinforced by the convergence of adaptive neuromodulation and implantable BCIs, which increasingly rely on the same technologies and face many of the same scientific, technical, and regulatory challenges [42]. These shared requirements highlight the need for a general-purpose, standardized translational ecosystem that enables accessible, reproducible, and scalable research across both fields [33], [41]. To support this vision, a translational ecosystem should satisfy several key design requirements. It should support both neural sensing and stimulation; provide an open research backend for integration into custom experimental protocols; and employ a modular and portable software architecture that supports multiple hardware platforms. Finally, the underlying technology should have a clear regulatory pathway to facilitate eventual clinical translation.

Guided by these principles, we developed an open translational ecosystem for adaptive neuromodulation and implantable BCI research by integrating CorTec’s Brain Interchange™ (BIC) device with the open-source BCI2000 framework [43]. The BIC is among the few investigational platforms that combine high-channel-count bidirectional neural interfacing, an open research interface, and a defined pathway for clinical investigation [33], [43]. While several software platforms support different aspects of neuromodulation research [33], including data acquisition [44], offline analysis [45], [46], [47], and real-time experimentation [48], [49], [50], [51], we selected BCI2000 because it is one of the most mature and widely adopted open frameworks, providing a comprehensive environment for complex adaptive neuromodulation and implantable BCI workflows. The resulting BIC-BCI2000 ecosystem establishes a common translational framework for the experimental paradigms presented in this work, enabling rapid implementation, evaluation, and transfer of protocols between other supported research hardware and BIC.

To support adoption of this ecosystem, we performed a comprehensive technical characterization and long-term validation of the BIC device. Although previous studies have reported selected aspects of the system [48], [51], [52], [53], [54], we independently verified and extended these evaluations while characterizing several capabilities that have not previously been comprehensively assessed.

The principal contribution is not a single application, but a common infrastructure through which investigators can implement neural sensing, functional mapping, biomarker detection, open-and closed-loop stimulation, evoked-potential recording, and BCI control. We additionally provide standardized characterization methods, software modules, imaging and analysis pipelines, experimental protocols, and openly accessible datasets. Together, these resources are intended to facilitate reproducible research and accelerate the evaluation of implantable neurotechnology across benchtop, preclinical, and clinical settings.

## 2. Materials & Methods

### 2.1. CorTec Brain Interchange–BCI2000 ecosystem

The BIC–BCI2000 ecosystem provides an open environment for rapid, iterative development and evaluation of implantable neuromodulation and brain-computer interface paradigms. To facilitate adoption, we provide comprehensive documentation, tutorials, and practical guidance for using the BIC–BCI2000 ecosystem through the BCI2000 website. The individual hardware and software components of the ecosystem are described below.

**CorTec’s BrainInterchange™** is a 32-channel implantable neural sensing and stimulation device capable of simultaneously acquiring signals from up to 32 channels sampled at 1 kHz. The system consists of three main components (Figure 1): (1) implantable unit responsible for neural recording, stimulation, and wireless data transmission; (2) an external unit comprising a communication unit, which facilitates communication between the implant and the computer, and a magnetically attached headpiece that inductively powers the implant; and (3) a personal computer, which processes the acquired signals and provides control of the system. The device is compatible with commercially available deep brain stimulation (DBS) electrodes as well as custom electrocorticography (ECoG) electrodes through standardized inline connectors [55], enabling the use of custom combined electrode configurations. Stimulation can be delivered through any subset of channels using asymmetric charge-balanced pulses (Supplementary Figure 4). A comprehensive description of the device architecture and capabilities is available in the manufacturer’s documentation, and previous publications [51], [52]. Importantly, the BIC software provides application programming interfaces (APIs) in multiple programming languages, enabling integration of the device’s full functionality into external research platforms such as BCI2000.

**Figure 1.**
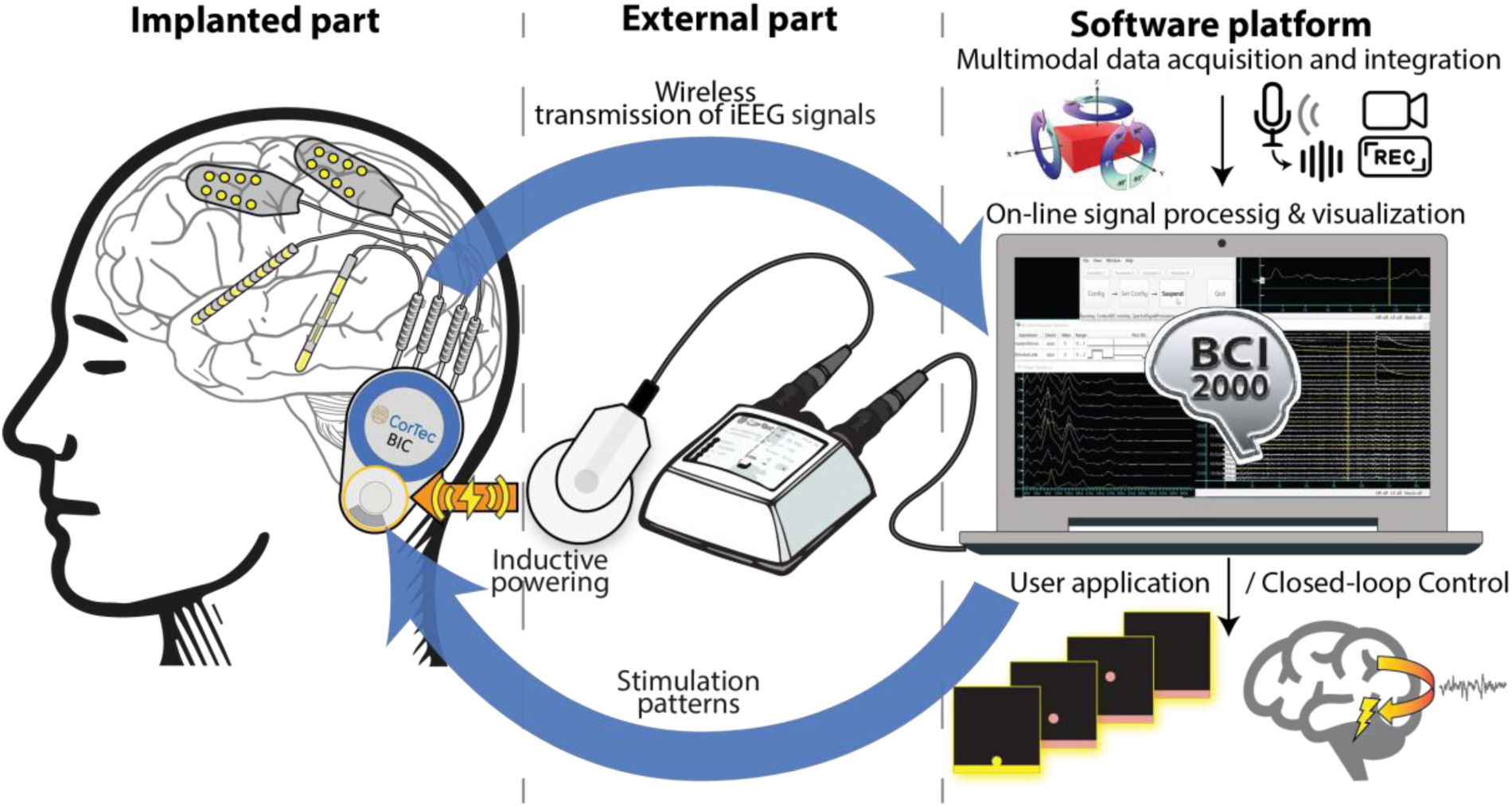
CorTec Brain Interchange (BIC) – BCI2000 ecosystem. The ecosystem consists of three parts: 1.) *Implantable part*: consisting of CorTec BIC device paired with cortical surface electrodes and/or deep-brain stimulation electrodes; 2.) *External part*: consisting of communication unit responsible for communication between the PC and implant and magnetically attached headpiece responsible for inductive powering of the implant; 3.) *Software platform*: handled by BCI2000 software responsible for integration, visualization and storage of multimodal recordings and on-line signal processing used to control the stimulation or user application.

**BCI2000** [56], [57] is a free, open-source (under GNU GPL), general-purpose software platform for adaptive neuromodulation and BCI research. Its modular, device-agnostic architecture enables rapid prototyping and facilitates the translation of established protocols across different hardware platforms. BCI2000 currently interfaces with more than 50 data acquisition platforms and a broad range of auxiliary stimulation and behavioral monitoring devices. Implemented in C++, BCI2000 provides a high-performance, low-latency, cross-platform framework for signal acquisition, processing, user feedback, and experimenter visualization. It also offers interfaces to high-level programming languages, including MATLAB, Python, Unity, and JavaScript, enabling custom signal processing, data analysis, and visualization. Another key feature of BCI2000 is its integrated acquisition and synchronization of multimodal data streams. By maintaining comprehensive annotations and metadata, the platform preserves data integrity and supports reproducible offline analysis. BCI2000 is a community-driven project with ongoing development and support coordinated by the National Center for Adaptive Neurotechnologies (http://www.bci2000.org).

### 2.2. In Vitro testing

We performed a series of in vitro experiments using the benchtop version of the BIC device (Figure 2.A) to comprehensively characterize its recording performance, including the amplifier noise floor, power-law fit parameters, transfer function, recording and stimulation latencies, power consumption, and impedance measurement accuracy. All benchtop recordings were conducted inside a custom-built Faraday cage to minimize environmental noise.

**Figure 2.**
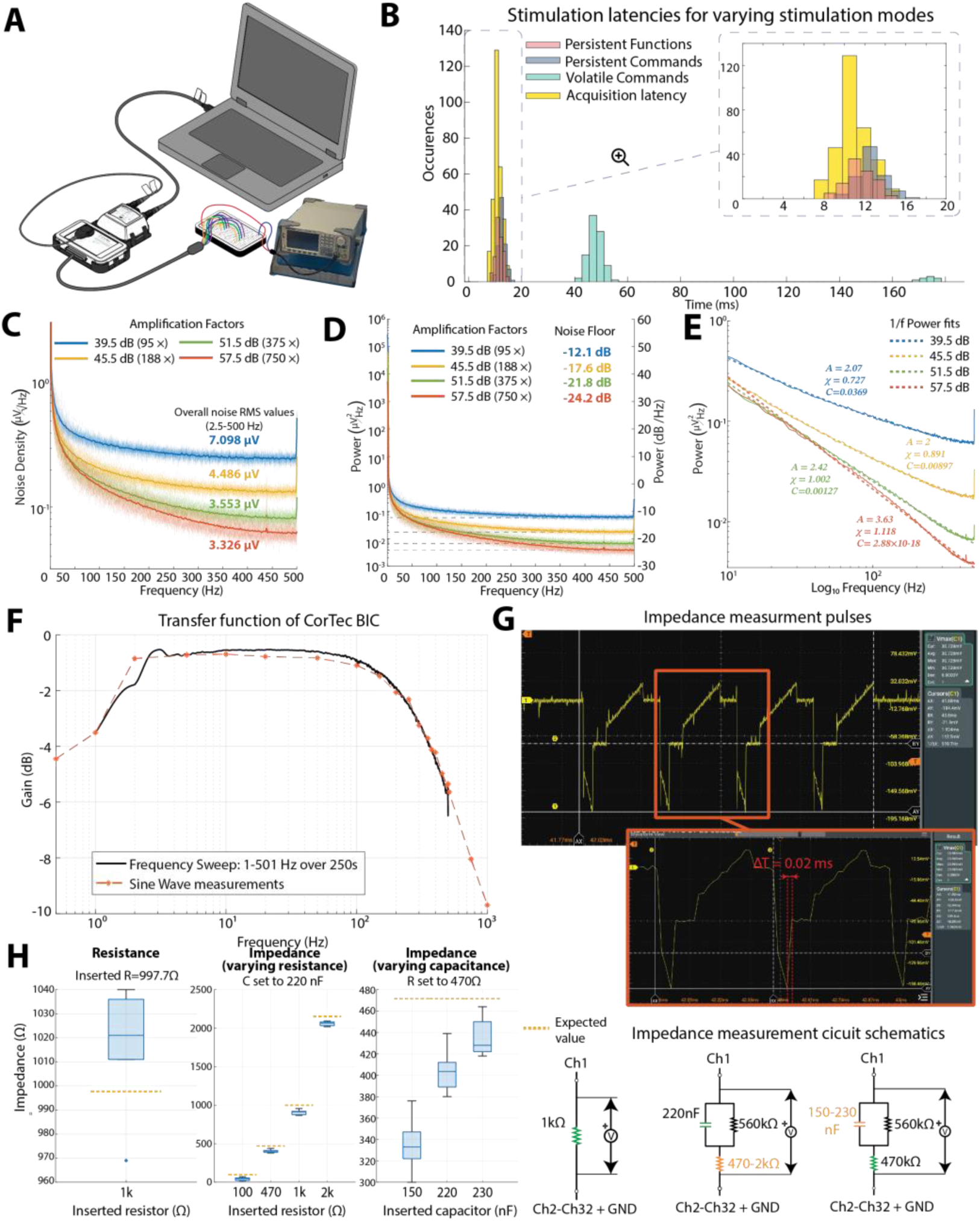
Benchtop testing. **A)** Recording setup for testing using benchtop version of CorTec Brain Interchange (BIC). **B)** Quantification of stimulation and recording latencies for various stimulation preloading modes: *Acquisition latency*: 10.89 ±1.59; *Persistent Function*: 12.68 ± 1.38; *Persistent Command*: 11.45 ± 1.19; *Volatile Commands*: 57.97 ± 34.45 ms. **C)** Amplifier root mean square (RMS) noise-density for different amplification factors. Noise density was calculated from a Welch periodogram of a shorted input recording within 2.5-500 Hz span at a 0.5 Hz frequency resolution. **D)** Power spectral densities of the shorted-input recordings and the corresponding noise floor (in dB) measured at 495 Hz. **E)** 1/f power fit for a shorted input periodogram calculated as in [58]. **F)** Transfer function of the CorTec BIC. Red asterisks mark the gain in decibels measured over 2 mV sinusoidal signal of specified frequency. The black line marks the gain calculated from peak-to-peak values of upper and lower envelope of the chirp signal. **G)** Zoomed-in impedance measurement pulse. The rise rime of ΔT was used to calculate effective frequency for impedance calculation. **H)** Results of impedance measurement tests. Impedance was measured over pure resistive load and combined resistive and capacitive load with varying resistance and capacitance. Results show that the device internal impedance measurement protocol is influenced mainly by the resistive load.

#### 2.2.1. Recording and stimulation latencies

As captured signal and stimulation commands are transmitted wirelessly between the BIC implant and the external hardware, it is important to quantify the associated system latency. Latencies were estimated by simultaneously operating two acquisition systems: the CorTec BIC, which performed neural recording and stimulation, and a low-latency reference device capable of generating transistor-transistor logic (TTL) pulses (g.USBamp; g.tec, Austria), which served as the timing reference. Upon a change in the experimental condition (BCI2000: StimulusCode), two events were triggered simultaneously: (1) generation of a TTL pulse by the reference device and (2) transmission of a stimulation command to the BIC. The TTL pulse served as the ground-truth reference against which both the recording and stimulation latencies of the BIC were measured. Wireless data from the BIC are streamed continuously every 2 ms, while online processing in BCI2000 is performed in user-configurable data blocks, introducing a buffering delay proportional to the selected block size (50 samples in this study). This buffering delay was excluded from the latency analysis. The BIC supports three stimulation modes, each with distinct advantages and limitations [48], [51]. Latencies were therefore evaluated separately for each preloading mode (Figure 2B).

#### 2.2.2. Amplifier noise floor

The amplifier noise floor was estimated from one-minute recordings acquired at each available amplifier gain setting. Power spectral densities (PSDs) were computed using MATLAB’s pwelch function with 2-second Hann windows and 90% overlap. Noise densities were then calculated as the root mean square (RMS) of the power spectra, corrected for the frequency-bin width of the spectral estimate.

#### 2.2.3. Power-law fits

The power-law fit parameters of the model in the form of *P* = *Af*^−**z**^ + *C* were estimated from the shorted input spectra by fitting the log-log least-squares as described in [58], over the frequency range 10-499Hz.

#### 2.2.4. Amplifier transfer function

The amplifier transfer function was estimated using two complementary approaches. First, a frequency sweep (chirp) generated by a signal generator was applied over the frequency range of 1– 501 Hz for 250 s with a constant peak-to-peak amplitude of 2 mV (Supplementary Figure 1). Second, 10 s pure sinusoidal signals were applied at discrete frequencies ranging from 0.1 to 1250 Hz for each amplifier gain setting. The transfer function was estimated from the chirp recordings by calculating the peak-to-peak amplitude of the smoothed signal envelope (solid black line in Figure 2F) and from the sinusoidal recordings by calculating the mean peak-to-peak amplitude across repeated measurements (dashed red line in Figure 2F). The measured peak-to-peak amplitudes were then compared with the input peak-to-peak amplitude, and the gain was expressed in decibels according to

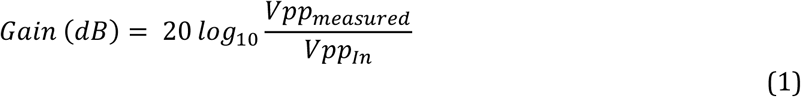

#### 2.2.5. Impedance measurement accuracy

The accuracy of the BIC impedance measurements was evaluated using repeated measurements (N = 10) of electrical circuits with known impedances (resistor tolerance: ±1%; capacitor tolerance: ±5%). The BIC estimates impedance by delivering a train of four short current pulses with an amplitude of 220 µA [51], [53], using the channel under test as the source and all remaining channels as the return. The resulting voltage response is averaged, and the impedance is calculated using Ohm’s law. To estimate the effective frequency of the impedance measurement, the probing pulses were captured using a laboratory oscilloscope (Rigol DHO1074), and the rise time (t_r_) was estimated from the falling edge of an individual pulse corresponding to the resistive discharge. The effective measurement frequency was then approximated as 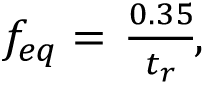, from which the corresponding angular frequency was calculated as *ω* = 2*πf_eq_*. Impedance measurements were then performed using both purely resistive loads and combined resistive-capacitive (RC) loads with varying resistance and capacitance values (Figure 2H). The theoretical impedance (yellow dashed line in Figure 2H) was calculated from the known circuit component values using the estimated angular frequency and the equations 2-5:

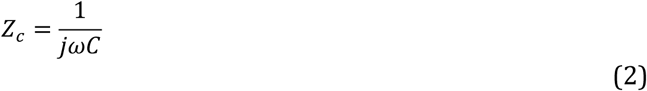

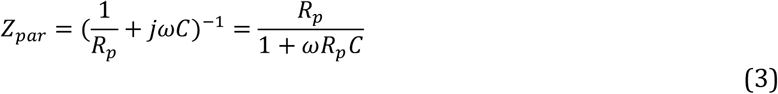

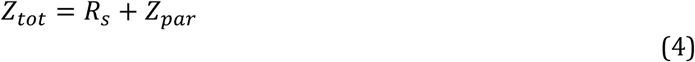

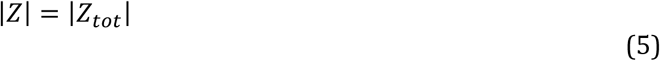

Where Z_C_ is the capacitor impedance, Z_par_ is the equivalent impedance of the parallel RC circuit, R_p_ is the resistance in the parallel circuit, and R_s_ is the resistance in series.

#### 2.2.6. In vitro stability testing

A 24-hour continuous recording was performed using the benchtop BIC unit placed inside a Faraday cage. Throughout the recording, power consumption, implant temperature, and packet loss (PL), defined as the proportion of data samples lost during wireless transmission, were continuously monitored (Supplementary Figure 5) using a USB voltage and current detection meter (FNIRSI FNB58 USB tester).

#### 2.2.7. Saline tests and hardware & software referencing strategy

Saline tests were performed on all implants before sterilization and implantation to verify device functionality and establish a reference baseline for recording and stimulation performance.

Measurements were conducted in a saline bath placed in a Faraday cage to minimize the environmental noise (Figure 3A). The testing protocol consisted of three recording conditions: (1) passive recordings to characterize the background noise, (2) active recordings of signals generated by a signal generator, and (3) stimulation recordings to verify the stimulation functionality of the implant. The recording configuration varied depending on the electrode leads available for each implant. Implants #1 and #3 used integrated ECoG electrodes, Implant #2 was tested using decommissioned DBS leads, and Implants #4 and #5 were evaluated using combined ECoG and DBS electrode configurations.

**Figure 3.**
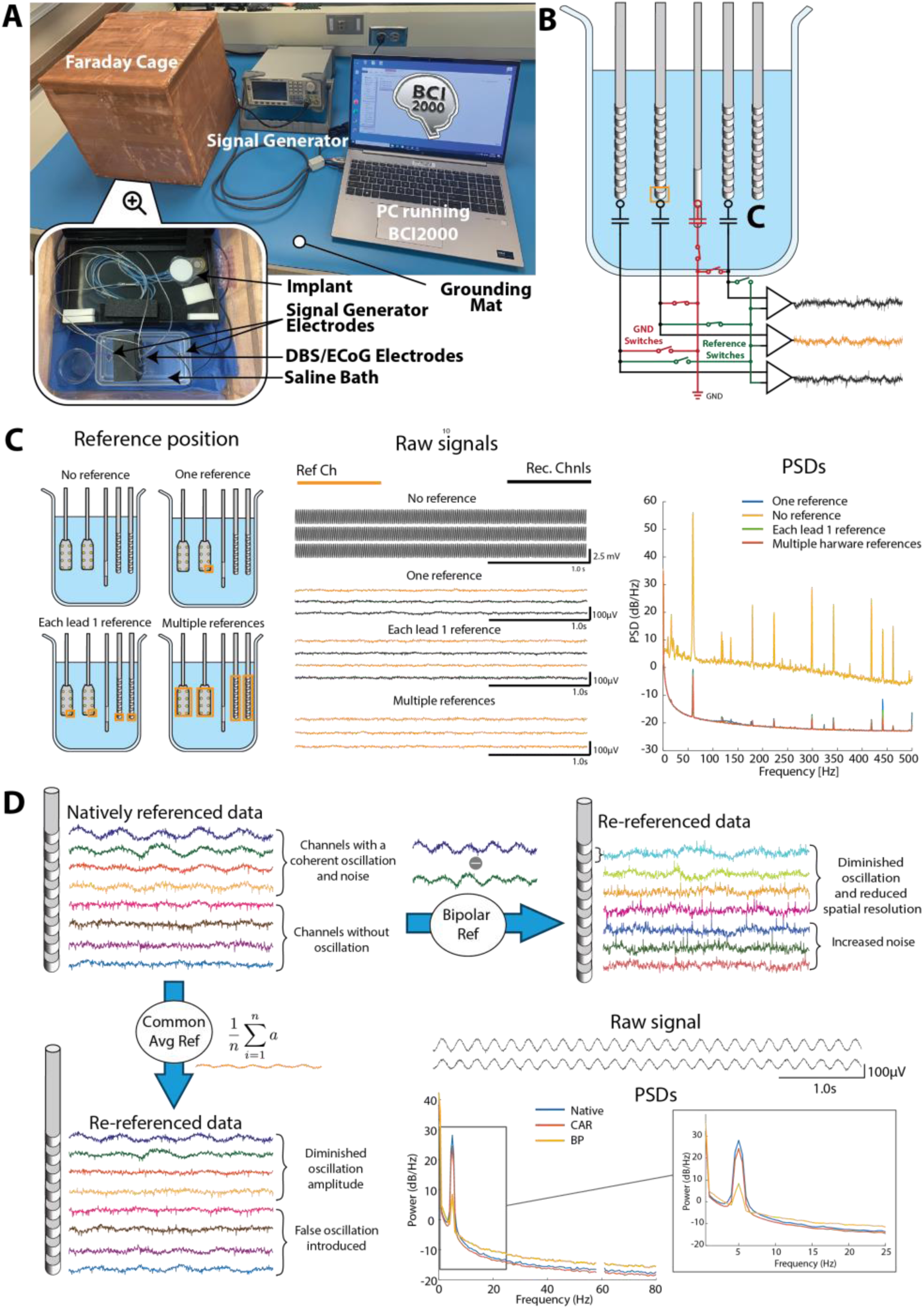
Saline test recordings. **A)** Recording setup. **B)** Schematic diagram of internal switching unit for specification of reference channel and utilization of ground electrode for recording in saline bath. **C)** Effect of hardware reference selection on quality of the signal and amplifier noise-floor. Selection of at least one reference channel is highly desirable. **D)** Influence of software re-referencing on signal quality and removal / induction of artifacts. Bipolar (BP) re-referencing is effective at removal of coherent noise across neighbouring channels at the cost of increasing the overall stochastic noise. Common average re-referencing (CAR) is effective at removing ergodic stochastic noise at the cost of possibility of inducing fake oscillations at other channels. The choice of the re-referencing method is therefore application dependent.

BIC supports dynamic selection of the hardware reference to one or multiple channels (Figure 3B). We performed a series of measurements to evaluate the effect of hardware reference selection on recording quality (Figure 3C and Supplementary Figure 6.) and to determine the optimal reference configuration for neural recordings.

In addition to hardware reference selection, we evaluated the influence of **software re-referencing** on signal quality and its ability to preserve different signal features. For this analysis, we used the single-reference saline recordings acquired under both passive and active recording conditions. Software re-referencing was performed using custom MATLAB functions for offline analysis and was implemented online using the BCI2000 Spatial Filter module.

### 2.2. In Vivo evaluation

To evaluate the readiness of the ecosystem for clinical translation, we conducted a series of preclinical studies in canine models (Canis familiaris) to assess the long-term stability and performance of the device, as well as to develop and validate experimental neuromodulation protocols. In addition, we performed a proof-of-concept study in a patient undergoing epilepsy monitoring (EMU) using the benchtop version of the BIC system connected to externalized stereoelectroencephalography (sEEG) leads.

### 2.3. Canine subjects and experiments

Five canines were implanted with the CorTec BIC system. Implant systems #2 and #4 were used in a canine with naturally occurring epilepsy, while the remaining implant systems were evaluated in neurologically healthy animals. All procedures were conducted under Mayo Clinic IACUC protocol A00001713-16-R19, and all animals were housed in a social environment. Canine models were selected because of their body size, which readily accommodates the implantable system, the occurrence of naturally arising neurological disorders analogous to those observed in humans, and their high degree of trainability. These characteristics enable neural recordings during translationally relevant behavioral paradigms in naturalistic environments. Furthermore, all canines are made available for adoption upon completion of the study^1^ allowing them to lead fulfilling lives beyond their participation in research. Details of the implanted animals and their corresponding electrode configurations are provided in *Table 1*.

**Table 1.** Implant subject information. Implant number, electrode configuration, total recorded time, number of days over which recordings were performed and average packet loss (PL) for each implant across all recordings.

| Implant # | Construct | Total recode time | Number of recording days | Average PL [%] ± STD |
| --- | --- | --- | --- | --- |
| Pivotal Implant (Laika) | ECoG (subdural) | 3h 45m | 17 | 4.05±1.66 |
| 1 (Belka) | ECoG (epidural) | 115h 50m | 489 | 3.50±2.73 |
| 2 (Billy) | DBS (HPC & ANT) | 23h 29m | 41 | 4.02±4.74 |
| 3 (Strelka) | ECoG (epidural) | 41h 11m | 173 | 3.24±4.64 |
| 4 (Willie) | Combined (ECoG + DBS) | 37h 48m | 91 | 3.93±4.01 |
| 5 (Mushka) | Combined (ECoG + DBS) | 2h 4m | 36 | 3.47±3.81 |
| Summary | - | 224h 7m | 847 | 3.56±3.63 |

ECoG– electrocorticography ; DBS – deep brain stimulation ; HPC – Hippocampus; ANT –Anterior nucleus of the thalamus;

#### 2.3.1. Surgical protocol

Canines were implanted with the BIC device. A schematic of the implant configuration is shown in Figure 4A. BIC was paired with either epidural ECoG electrode strips or DBS leads (Boston Scientific Cartesia) (Figure 4B). The implantable device was placed subcutaneously over the torso, with the electrode leads tunneled subcutaneously to the cranial implantation site. The surgical procedure has been described previously (Schalk et al., 2022 [43]), and an updated version of the protocol is available online at https://wustl.app.box.com/s/hq5rgw4mnb70nd6f07jkk50u2o70n46s.

**Figure 4.**
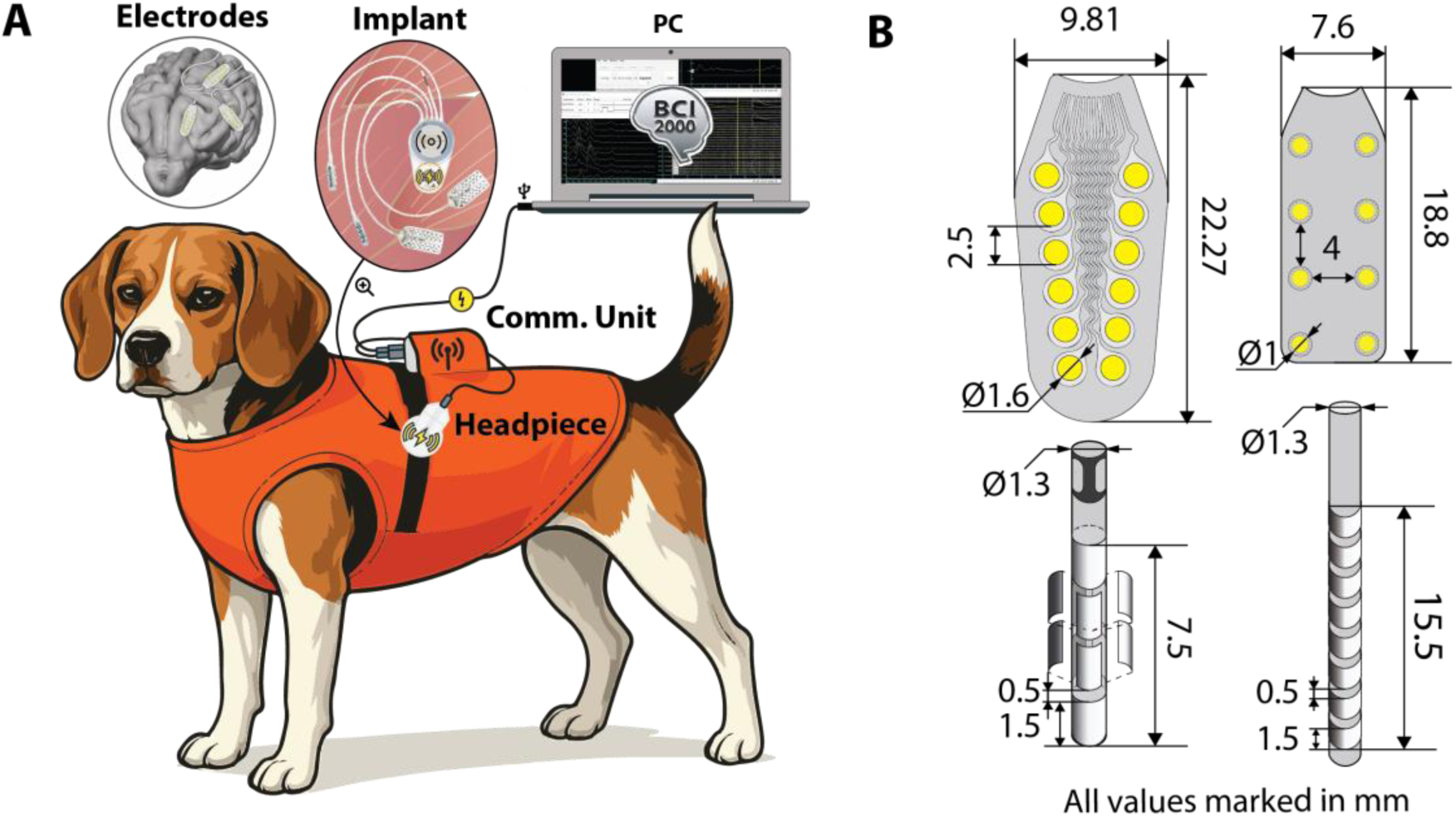
CorTec BrainInterchange – BCI2000 recording setup in canines. All animal procedures were approved under Mayo Clinic IACUC protocol A00001713-16-R19. The canines will be made available for adoption upon completion of the study†. **A)** The Brain Interchange (BIC) device is implanted in the scapular region, with electrode leads tunnelled subcutaneously to the cranial implantation site. During recording sessions, the canines are trained to remain stationary and are not physically restrained. A calming vest is worn to provide additional stabilization of the magnetically attached external headpiece and to house the communication unit within a dedicated pocket. **B)** Dimensions and configurations of the electrode interfaces compatible with the BIC system, including cortical paddle electrodes, and directional or sequential deep brain stimulation leads. † In compliance with the Minnesota state Beagle Freedom Bill statute 135A.191.

#### 2.3.2. Electrode localization and anatomical co-registration

Each subject underwent preoperative 3T magnetic resonance imaging (MRI) for surgical planning.

Following implantation, a postoperative computed tomography (CT) scan was acquired, and additional follow-up CT scans were obtained several months later to assess potential electrode displacement. All imaging data were processed using a custom image-processing pipeline. MRI and CT volumes were first reoriented to the RAS coordinate system and subsequently co-registered using SPM [59]. Brain tissue was semi-automatically segmented from the preoperative MRI using tissue probability maps derived from the stereotaxic breed-averaged canine brain atlas [60] in SPM [59].

The resulting segmentations were manually refined in 3D Slicer [61], [62], [63], [64]. The co-registered CT images and brain masks were then used to skull-strip the preoperative MRI. The skull-stripped MRI was nonlinearly normalized to the Stereotactic Cortical Atlas of the Domestic Canine Brain from Johnson et al. [65] using ANTs [66], [67], [68], which subsequently served as the template space for cortical anatomical labelling. To improve the localization of subcortical structures, we generated a custom atlas by combining the Johnson atlas with the Detailed Canine Brain Label Map for Neuroimaging Analysis [69], with the Johnson atlas serving as the common template space.

Additional anatomical structures were manually incorporated from The Brain of the Dog atlas [70] using 3D Slicer. Electrode locations were determined from the co-registered postoperative CT. ECoG electrodes were localized using the CTMR package [71] as previously described [72]. DBS leads were localized using Lead-DBS [73] [74] and DBS reslice [75]. The same software packages were used to generate three-dimensional brain renders and electrode visualization.

#### 2.3.3. Longitudinal recordings

Longitudinal recordings were acquired from each canine to evaluate the long-term stability and performance of the implanted system. Recordings were performed on regular basis (approximately 3 times a week) and consisted of approximately 3-minute passive recordings while the animals were resting calmly. Impedance measurements were acquired during the same recording sessions. The recordings were analyzed using a custom MATLAB processing pipeline. For each recording, the recording duration, PL, and the number of bad channels were extracted. Spatial filtering was then applied using each re-referencing strategy, after which temporal signal statistics (variance and RMS) and spectral statistics (band power and PSD RMS) were computed. These metrics were used to evaluate system performance and determine the optimal spatial filtering strategy for signal preprocessing.

#### 2.3.4. Automatic bad channel detection

Bad channels were identified from the longitudinal recordings using the following algorithm:

1) Prior exclusion: Channels with known defects were excluded before analysis based on prior information, including pre-implantation saline recordings or postoperative imaging. In addition, channels with an impedance greater than 10 kΩ were automatically classified as bad when impedance measurements were available.
2) Candidate selection: Candidate bad channels were identified using two complementary criteria:

a) Time-domain analysis: Channel-wise signal standard deviation was evaluated to identify statistical outliers.
b) Spectral analysis: Outliers were identified by comparing the signal power within the 1–150 Hz band and the high-frequency noise floor, estimated over the frequency range from 0.75 × the Nyquist frequency to the Nyquist frequency.
3) Candidate aggregation: Candidate channels identified by the individual metrics were combined using robust statistics, including lead-wise threshold refinement and integration of the time-domain, spectral, and lead-based detection criteria.

#### 2.3.5. Functional mapping

Functional mapping experiments were performed in two canines using four task paradigms: simple visual, visual flicker, auditory, and motor. Each task was repeated across multiple runs until a sufficient number of active and inactive task blocks were acquired (78–259 blocks, depending on the task). All experiments were conducted with the animals unrestrained in a quiet room.

- **Simple visual task**: The room lights were turned off, and the animal was positioned in front of a computer monitor. The visual stimulus consisted of a blank white screen presented for 5 s.
- **Visual flicker task**: The room lights were turned off, and the animal was positioned in front of a computer monitor. The stimulus consisted of a blank white screen flickering at 5 Hz (100 ms on, 100 ms off) for 2 s.
- **Auditory task**: The animal was positioned next to a computer that periodically presented a 440 Hz pure tone for 5 s.
- **Motor task**: The animal was positioned in front of the examiner, who received timing cues from the computer and periodically moved the forepaw contralateral to the implanted hemisphere. Limb movement was recorded using an Xsens accelerometer attached to the paw with a Velcro strap and was manually segmented during offline analysis.

Collected data were then post-hoc analyzed in MATLAB 2023b (MathWorks, Inc., Natick, MA, US). Bad channels were excluded from the analysis. Signals were high-pass filtered using a third-order Butterworth filter with a cutoff frequency of 0.5 Hz to remove the DC offset and subsequently re-referenced using the lead common average (LeadCAR) method, in which the average voltage across all non-bad channels within each lead was subtracted from every channel in that lead. PSDs were then calculated for each channel and task using 1 s Hann windows with 50% overlap and normalized to the

PSD computed over the entire recording. Task blocks were visually inspected, and trials containing artifacts were excluded from further analysis. The normalized PSDs were subsequently compared across task conditions using the r^2^ metric as described in our previous work [72], [76]. Finally, changes in low-frequency power (task dependent) and high-frequency power (65–175 Hz, excluding line-noise harmonics) were projected onto the rendered brain for visualization as described in [72], [77] and [76].

#### 2.3.6. Closed-loop stimulation

Closed-loop stimulation experiments were performed in Implant #4 (Willie) while the animal was sleeping in a quiet room. Stimulation was delivered through the thalamic DBS lead. Recording channels were selected during the experiment by visually inspecting the online spectral estimates of neural activity during sleep. Based on this assessment, two bipolar derivations exhibiting distinct spectral peaks were selected for closed-loop control.

Closed-loop stimulation was implemented using a custom processing pipeline built from BCI2000 modules (see Supplementary Figure 14 for a detailed description of the algorithm). Stimulation was triggered when the control signal exceeded a user-defined threshold and remained above that threshold for a specified duration (0.5 s in this study). The control signal was derived from the z-normalized spectral power computed over a 30 s normalization window in the 20–24 Hz and 75–85 Hz frequency bands from two bipolar derivations (Channels 9–13 and 30–31). The 1–100 Hz spectral power was estimated using an autoregressive model [78] applied to 1 s Hann-windowed data. In addition to this implementation, alternative closed-loop pipelines were developed in which the control signal was derived from the time-domain signal envelope or from spectral power normalized to a baseline period, analogous to the normalization used for the r^2^ analysis. To ensure stimulation safety and prevent self-triggering, the algorithm incorporated a user-defined refractory period of 2.5 s following each stimulation pulse, during which the recorded signal was excluded from online analysis.

#### 2.3.7. Recording and modulation of epilepsy biomarkers

Implants #2 and #4 were evaluated in a canine with naturally occurring epilepsy. Interictal epileptiform discharges (IEDs) were detected in both longitudinal recordings and long-term stimulation recordings using a previously established spike detection and artifact rejection algorithm [79], [80], [81]. For both implant systems, chronic stimulation was delivered for 30 min per day over a period of 6 weeks, divided into three 10-minute stimulation sessions. However, only Implant #4 included baseline recordings acquired using the same recording parameters as the stimulation sessions, enabling a direct comparison between baseline and stimulation conditions. Differences in IED rates between baseline and stimulation conditions were evaluated using a generalized linear model (GLM) with a negative binomial distribution to account for overdispersion in the spike count data. Raw spike counts were modeled as the response variable, with stimulation condition as the primary predictor and baseline serving as the reference category. To account for differences in recording duration and the number of analyzed channels, the logarithm of the recording exposure (recording duration × number of valid channels) was included as an offset, thereby modeling IED rates per channel-minute. Model coefficients were exponentiated to obtain incidence rate ratios (IRRs) with 95% confidence intervals (95% CI), representing the relative change in IED rate relative to baseline. Statistical significance was assessed using two-sided Wald tests, with p < 0.05 considered statistically significant.

#### 2.3.8. Single-pulse electrical stimulation

Single-pulse electrical stimulation (SPES) experiments were performed using Implant #5. Stimulation was delivered through the thalamic lead (Boston Scientific Cartesia directional lead), and evoked responses were recorded from the cortical electrodes. All recordings were acquired using identical stimulation parameters: stimulation amplitude of 3 mA, primary pulse width of 200 µs, dead zone 0 of 10 µs, dead zone 1 of 10 µs, and an inter-pulse interval of 3 s. Recording parameters included an amplifier gain of 45.5 dB, no specified reference channel and the use of a ground electrode during recording. The raw recordings were re-referenced using nearest-neighbor bipolar derivations and segmented into 2 s epochs spanning −0.5 to 1.5 s relative to the stimulation onset. Evoked responses were then quantified using the canonical response parameterization method described by Miller et al. [82]. Example responses were selected for visualization and projected onto the rendered brain surface to facilitate anatomical interpretation of the results.

### 2.4. Human proof-of-concept evaluation

One human participant (male, 34 years) was enrolled in this study following implantation of sEEG electrodes as part of his clinical evaluation for drug-resistant epilepsy. Electrode placement was determined solely by clinical considerations and was not modified for research purposes. The participant provided informed consent to participate in the study and consent for audio and video recording for publication purposes. The experiment was conducted in the adult EMU at Mayo Clinic, Rochester, Minnesota, and the study was approved by the Institutional Review Board of the Mayo Clinic (IRB 15-006 530).

#### 2.4.1. Brain-computer interface cursor control

The human participant completed a motor screening task followed by one-dimensional cursor control using the BCI2000 platform and a g.tec g.HIamp amplifier, as previously described by Jensen et al. [83]. Briefly, the motor screening consisted of cued hand, tongue, and foot movements, together with kinesthetic motor imagery, and was used to identify the bipolar derivation exhibiting the strongest task-related modulation (highest r^2^ value). After functional mapping, signal acquisition was switched from the g.HIamp to the CorTec BIC while using the corresponding bipolar derivation identified during functional mapping for real-time BCI cursor control.

Cursor control was based on high-frequency activity (72.5–112.5 Hz) extracted from the selected bipolar derivation using the standard BCI2000 real-time signal processing pipeline. During each trial, the participant performed either overt hand movement or kinesthetic motor imagery, depending on the target location, to control the cursor toward the displayed target. Further details of the experimental protocol, signal processing, decoder calibration, and performance evaluation are provided in the prior work by Jensen et al. [83].

## 3. Results

### 3.1. In vitro evaluation

Benchtop test recordings (Figure 2A) first focused on characterizing the wireless transfer latencies of the BIC system. The measured **acquisition latency** was 10.89 ±1.59 ms. **Stimulation latency**, defined as the interval between issuing the stimulation command and the onset of stimulation, depended on the selected preloading mode (Figure 2B). The persistent function mode exhibited a latency of 12.68 ±1.38 ms, while the persistent command mode achieved the lowest latency of 11.45 ± 1.19 ms. In contrast, the volatile command mode showed substantially higher latency (57.97 ± 34.46 ms) and a bimodal distribution, with a dominant peak centered at approximately 47 ms and a secondary peak around 175 ms. The second peak likely reflects delayed command delivery caused by retransmission when the initial communication attempt is unsuccessful.

Having established the timing characteristics of the system, we next characterized its intrinsic recording performance. The **overall RMS noise** progressively decreased with increasing amplifier gain (Figure 2C), from 7.098 µV at 39.5 dB to 4.486 µV, 3.553 µV, and 3.326 µV at amplification factors of 45.5, 51.5, and 57.5 dB, respectively. Corresponding **PSDs** (Figure 2.D) revealed that the shorted-input spectra exhibited 1/f-like behavior at low frequencies before transitioning to a flat noise floor at approximately 200 Hz, 250 Hz, 350 Hz, and beyond 400 Hz for amplification factors of 39.5, 45.5, 51.5, and 57.5 dB. **Power-law fits** confirmed the characteristic 1/f behavior of the amplifier noise. As the amplifier gain increased, the reduction in the white-noise floor extended the frequency range over which the spectra followed the power-law relationship, resulting in progressively larger fitted values of the power-law exponent χ (Figure 2E).

To determine the practical recording bandwidth of the system, we next evaluated its frequency response. The **transfer functions** estimated from the frequency sweep (Supplementary Figure 1) and discrete sinusoidal measurements were in close agreement (Figure 2F). Both methods demonstrated a nearly flat frequency response between 2 and 100 Hz, with the gain decreasing below −3 dB at approximately 1 Hz and 300 Hz. These measurements are consistent with the manufacturer’s specified hardware passband of 2–325 Hz.

Following characterization of the recording performance, we evaluated the accuracy of the built-in impedance measurement. The BIC estimates **impedance** using a train of four probing current pulses (Figure 2G), which were captured using a laboratory oscilloscope. The rise time of the resistive component of the stimulation pulse was estimated to be approximately 0.02 ms (Figure 2G), corresponding to an effective measurement frequency of 17.5 kHz. This frequency was subsequently used to calculate the theoretical impedance of the test circuits. For a theoretical impedance of 100 Ω, the measured impedance was 46 Ω (IQR: 23–55 Ω), whereas a theoretical impedance of 2152 Ω was measured as 2066 Ω (IQR: 2033–2081 Ω) (Figure 2H). Similarly, increasing the capacitance while maintaining the same resistive load resulted in progressively lower measured impedances than predicted by calculations. The impedance measurements for purely resistive loads yielded a median BIC impedance of 1021 Ω for a 997.7 Ω resistor measured with a multimeter (IQR: 1011–1036 Ω).

We finally assessed the long-term stability of the benchtop system during continuous operation. A **24-hour benchtop recording** was performed to characterize the long-term power consumption of the BIC system (Supplementary Figure 5). After an initial transient, the power consumption of the whole system from the source stabilized at approximately 1.32 W and remained constant throughout the recording. Implant temperature remained below 30.5 °C and the average PL was 0.8%, confirming stable system operation.

In addition to the benchtop characterization, we evaluated the functionality of each implant under saline conditions (Figure 3A). **Saline measurements** confirmed the functionality of all implants prior to implantation and identified two faulty channels in Implant #3. These experiments also enabled evaluation of the effects of hardware reference selection, grounding, and software re-referencing on recording quality (Figure 3B). Hardware reference selection substantially reduced the overall noise floor compared with recordings acquired without a hardware reference (Figure 3C). Using at least one reference electrode markedly reduced background noise, whereas employing multiple reference electrodes produced little additional improvement. The position of the selected reference electrode also influenced the amplitude of recorded oscillatory activity (Supplementary Figure 6). Software re-referencing further altered the signal characteristics depending on the selected spatial filtering strategy (Figure 3D).

### 3.2. In vivo evaluation

A total of **six BIC devices were implanted across five canines**. The canine implantation setup is illustrated in Figure 4A. The BIC was implanted with either epidural ECoG strips or DBS leads (Figure 4B and Figure 5). The initial pivotal implantation, performed in 2021 using an experimental version of the BIC system, was reported previously by Schalk et al. [43]. The subsequent five implant systems employed the current-generation BIC device and were followed longitudinally for a total of 2,296 implant-days (mean 459 ± 438 days per implant), with the longest follow-up exceeding three years (Figure 6A). Implant #2 was explanted approximately three months after implantation because of an antibiotic-resistant infection. Following complete recovery, the same animal was successfully re-implanted with Implant #4.

**Figure 5.**
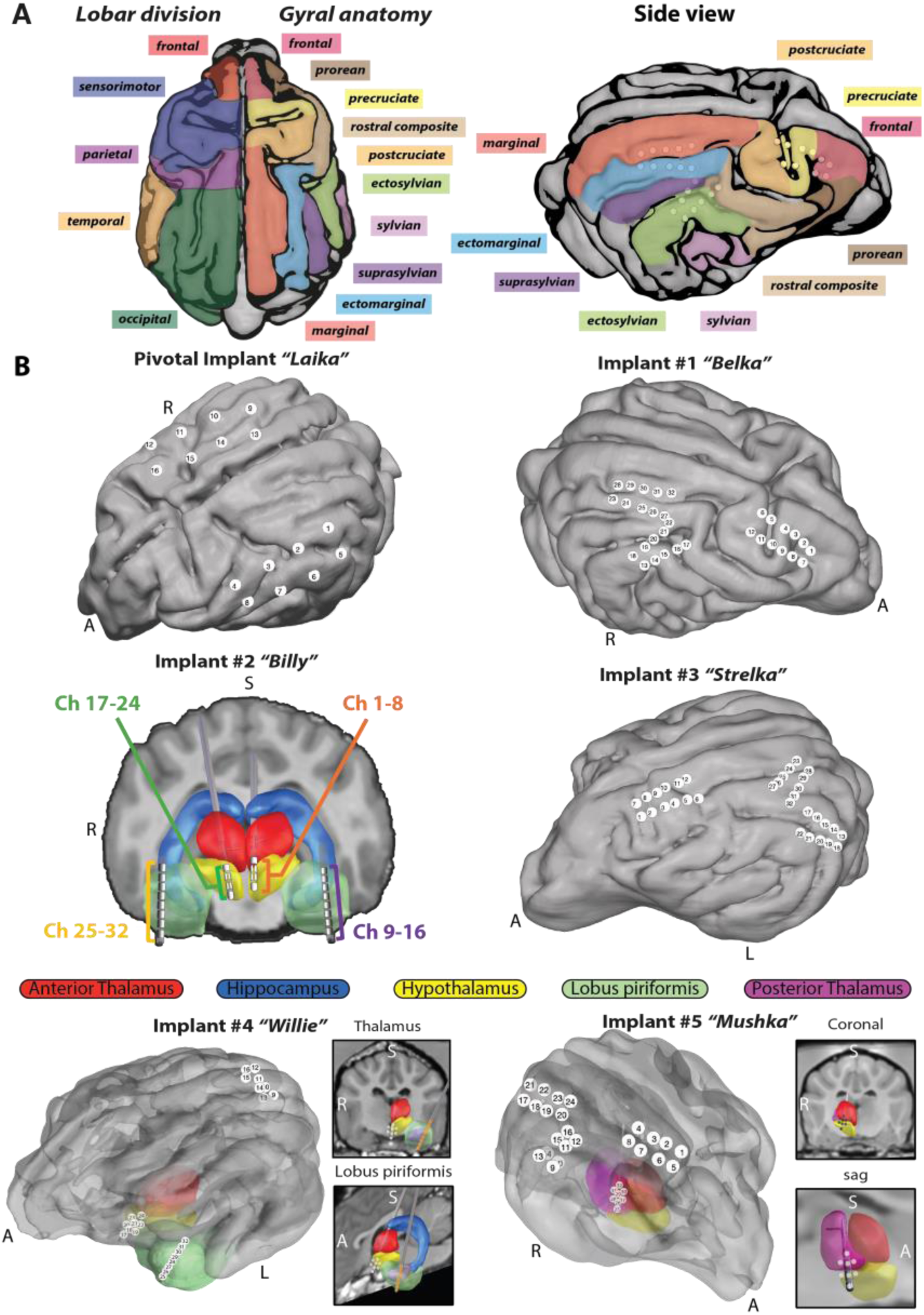
Electrode locations. **A)** Lobar and gyral anatomical parcellation of the canine brain based on the canine brain atlas developed by Johnson et al. [65]. **B)** Electrode locations projected onto subject-specific brain surface renderings. Electrode coverage targeted motor, temporal, and visual cortical regions, as well as selected deep brain structures. Surface electrodes were implanted epidurally through a cranial opening, while depth electrodes were placed stereotactically within the targeted subcortical structures.

**Figure 6.**
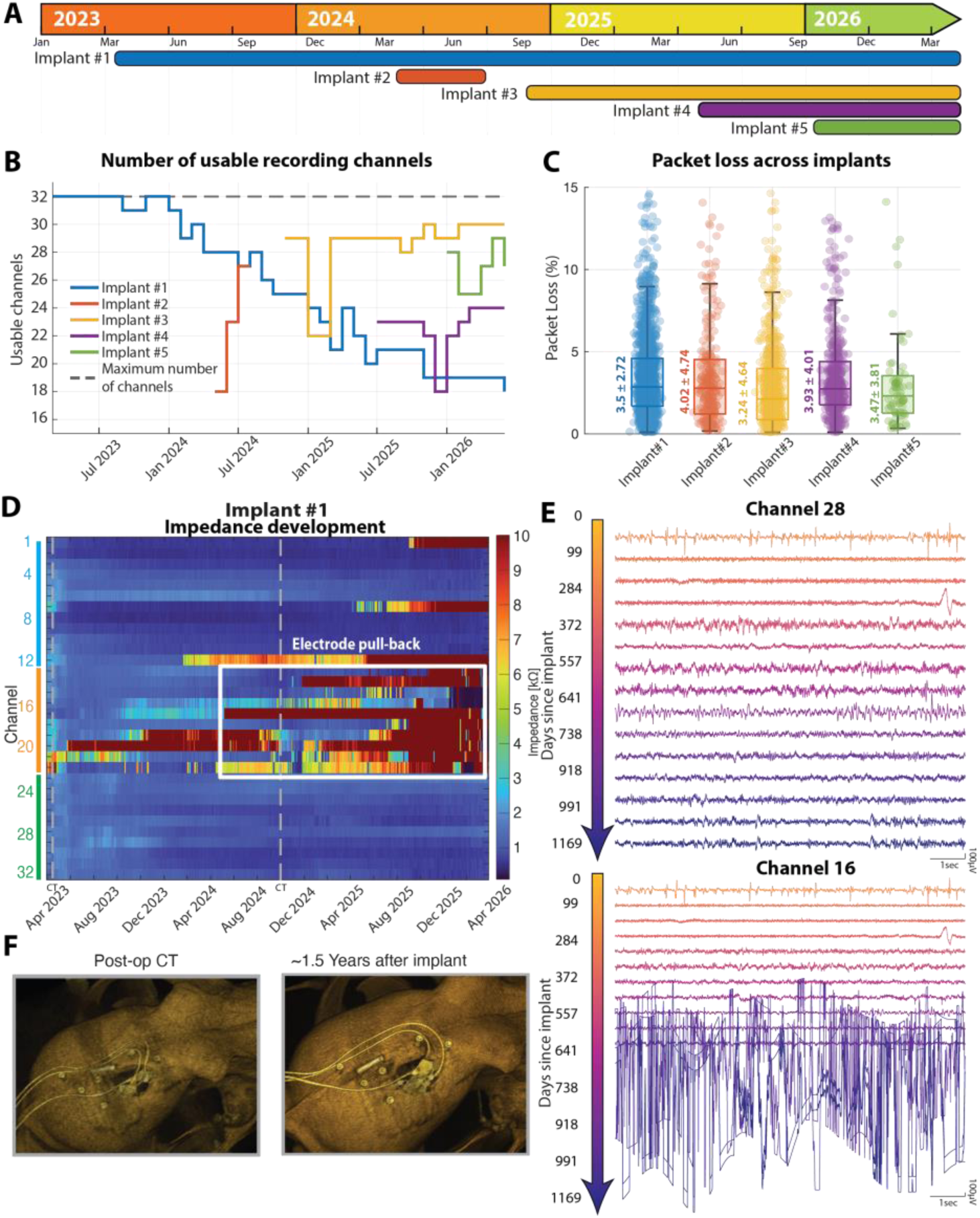
BIC–BCI2000 long-term recording capability. **A)** Implantation timeline for all canine subjects. **B)** Number of usable recording channels over time for each implant. Channel usability was determined using an automated quality assessment algorithm, allowing channels to transition between usable and unusable states throughout the implantation period. **C)** Distribution of wireless data packet loss across implants. The middle line indicates the median, and the adjacent text reports the mean ± standard deviation. **D)** Longitudinal electrode impedance measurements for Implant #1, the longest implanted device. Although some channel failures were attributable to electrical malfunction, the majority were associated with mechanical failure resulting from increased stress on the implanted leads during unrestricted movement of the canines. **E)** Representative long-term signal evolution of two recording channels over more than three years: one channel that remained stable throughout the implantation period and one channel that exhibited gradual signal degradation. **F)** Computed tomography (CT) renderings of Implant #1 demonstrating mechanical displacement of implanted electrodes.

**Longitudinal passive recordings and impedance measurements** were acquired throughout the follow-up period to evaluate the long-term recording stability of the implanted systems. Based on these recordings, the quality of individual recording channels was assessed automatically, and the number of functional channels was tracked over time (Figure 6B). Implant #1 exhibited a gradual decline in the number of functional recording channels, whereas Implants #2, #3, and #4 showed stable or improved channel availability during follow-up. Across all implants and recording sessions, the average PL remained below 5% (Figure 6C), indicating stable long-term wireless communication.

To better understand the observed channel failures, the longitudinal recordings were evaluated together with impedance measurements and follow-up CT imaging. Most channel failures were associated with mechanical changes at the electrode interface, as evidenced by progressive impedance changes (Figure 6D and Supplementary Figure 8), corresponding deterioration in signal quality (Figure 6E, bottom), and postoperative CT imaging demonstrating electrode displacement in Implant #1 (Figure 6F). In contrast, channels unaffected by mechanical or electrical failures maintained stable recording quality throughout the implantation period (Figure 6E, top).

**Functional mapping experiments** were performed approximately two years after implantation in Implant #1 and one year after implantation in Implant #3, demonstrating sufficient long-term recording stability to reliably capture task-related cortical activity. Analysis of the functional mapping recordings revealed task-related broadband activation (65–175 Hz) across both animals (Figure 7).

**Figure 7.**
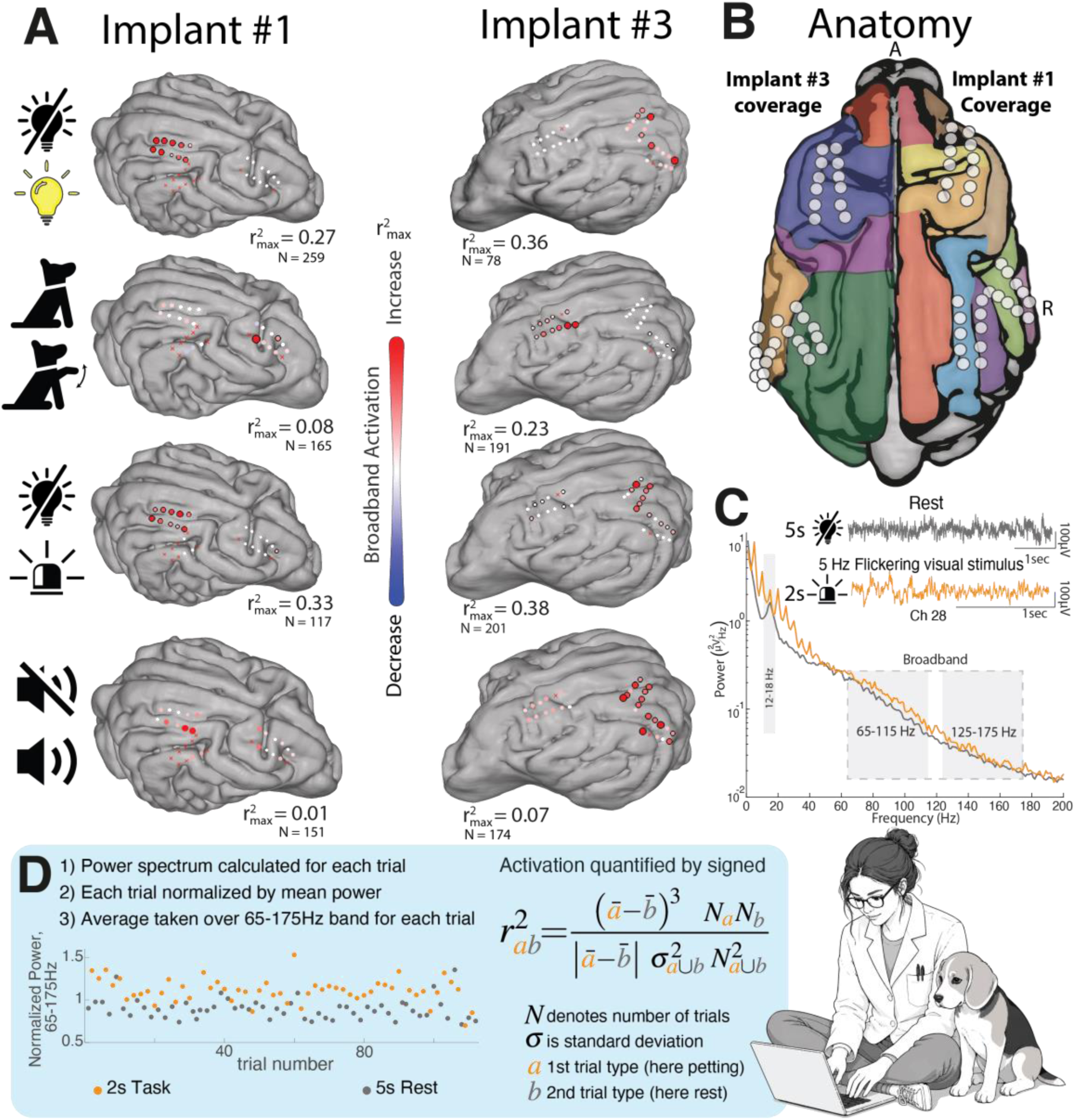
Functional mapping using block-design tasks. **A)** Broadband (65–175 Hz) activation maps for individual sensory tasks in two canines, scaled by the maximum coefficient of determination (r²). Visual stimulation produced the strongest activation in the occipital cortex, whereas motor tasks elicited activation primarily within the frontal cortex, centered around the rostral composite and precruciate gyri. No substantial task-related activation was observed during the auditory paradigm within the implanted electrode coverage. **B)** Canine cortical anatomy, color-coded as in Figure 5, with the implanted electrode locations overlaid. **C)** Representative signals and corresponding power spectral densities (PSDs) from a single trial of the visual flicker stimulation task. Notice the stimuli induced artifact present at 5Hz and its harmonics. **D)** Schematic illustrating the calculation of the r² activation coefficient used to quantify task-related neural activation.

The strongest responses were observed during the visual paradigms. Both the simple visual and 5 Hz flickering visual stimulation elicited robust broadband activation (r^2^ = 0.27–0.38) localized primarily to the ectomarginal and suprasylvian gyri within the occipital cortex, with additional activation extending into the posterior ectosylvian gyrus of the temporal cortex (Figure 7A,B). In addition to broadband activation, the flickering visual stimulus evoked a pronounced steady-state visual evoked potential (SSVEP) at the stimulation frequency (Figure 7C). The motor task (contralateral forepaw movement) produced spatially confined broadband activation over the precruciate and postcruciate gyri within the sensorimotor cortex. Although the maximum response observed in Implant #1 was relatively modest (r^2^ = 0.08), the activation remained highly localized. In contrast, the auditory task elicited only weak broadband modulation (r^2^ = 0.01–0.07), and the activated regions were not spatially consistent between the two animals.

**Closed-loop stimulation** was implemented and evaluated during sleep recordings in Implant #4 (a canine with naturally occurring epilepsy). During sleep, prominent oscillatory activity was observed in two distinct recording sites: 20–24 Hz oscillations in the piriform lobe and narrowband high-gamma activity (75–85 Hz) in the cortex. These spectral features were used independently as control signals to trigger low-amplitude thalamic stimulation (250 µA). Figure 8 illustrates the recording and stimulation sites, with the cortical recording channel highlighted in green and the piriform recording channel in orange. The online processing pipeline detected sustained increases in spectral power and generated stimulation whenever the normalized control signal exceeded the predefined threshold (2.0 for the cortical recording site and 2.5 for the piriform recording site). The detector selectively responded to sustained increases in spectral power while ignoring transient fluctuations, resulting in robust closed-loop triggering across both recording sites. Sustained increases in power within the selected frequency band preceded stimulation onset by approximately 1.5 s. This delay was determined by the temporal integration window and persistence criteria implemented in the detection algorithm.

**Figure 8.**
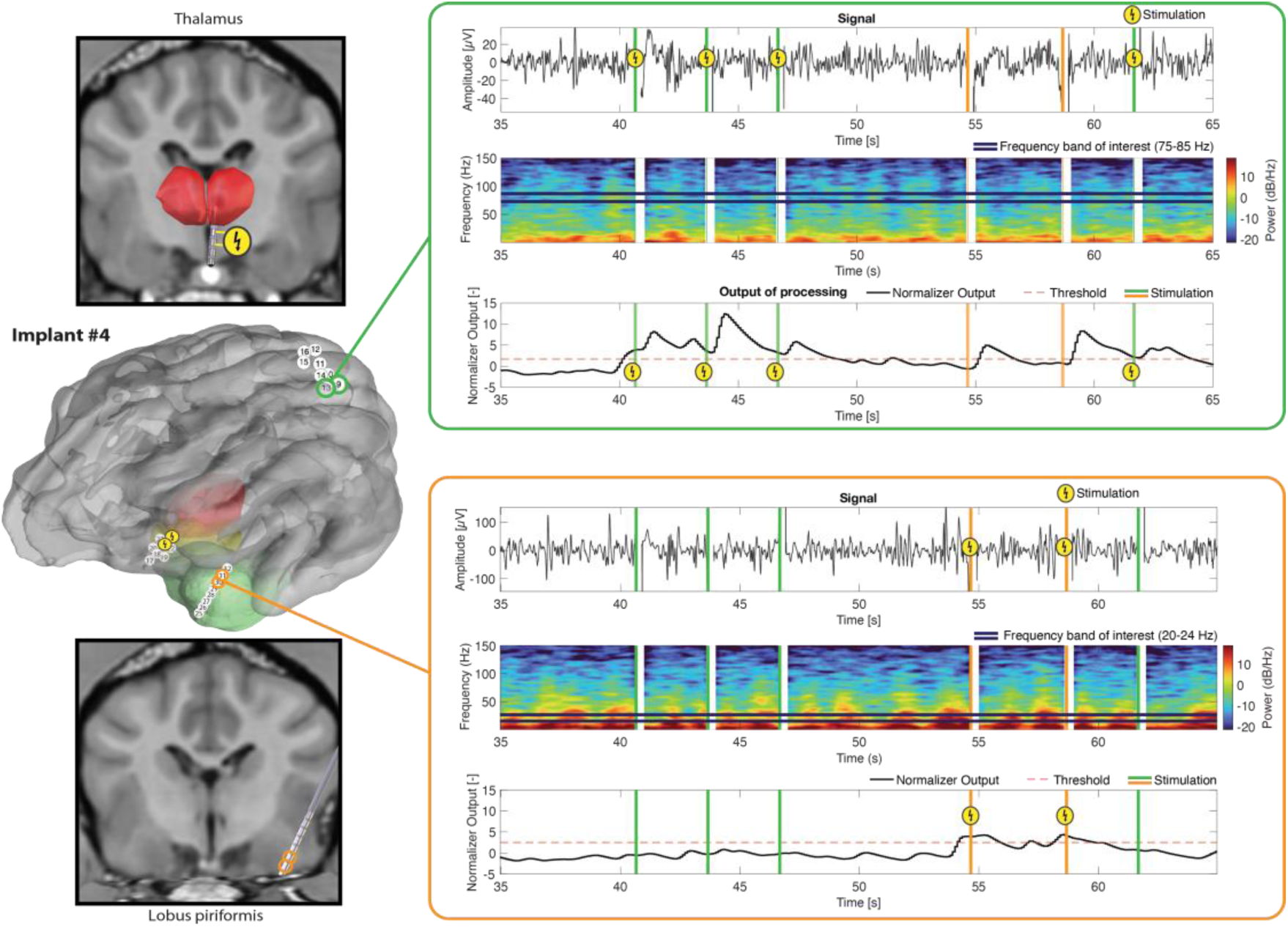
Closed-loop stimulation triggered by online detection of neural activity. **Left:** Two bipolar electrode pairs were selected for neural sensing and real-time signal processing. The sensing locations were arbitrarily chosen and are indicated by the orange and green circles. Stimulation was delivered through a thalamic depth electrode, indicated by the yellow circle with a lightning bolt. **Right:** Representative recordings from the sensing channels, corresponding time-frequency representations, and outputs of the online signal processing pipeline. Signal processing consisted of bipolar re-referencing, real-time spectral estimation using 1-second analysis windows, extraction of power within a control frequency band (18–24 Hz or 75–85 Hz), and z-score normalization relative to the preceding 30 seconds of recording while excluding periods containing stimulation. Stimulation was triggered when the normalized signal exceeded a predefined threshold. In this example, the latency between the onset of the neural event and stimulation delivery was approximately 1.5 seconds. This latency was primarily determined by the spectral analysis window length and other signal processing parameters and can be adjusted according to the requirements of the closed-loop application.

Longitudinal recordings were obtained from a **canine with naturally occurring epilepsy** using implants #2 and #4 (Figure 9A). IEDs were consistently detected throughout the implantation period, demonstrating stable long-term recording of epileptiform biomarkers (Figure 9B and Supplementary Figure 15). In Implant #4, we additionally captured an electrographic seizure that was independently determined by a clinical epileptologist (GAW) to originate from the seizure onset zone (SOZ) located at the bipolar derivation between channels 25–26 within the piriform lobe (Figure 9C). Implant #4 was subsequently used to evaluate the effects of chronic daily stimulation on IED incidence. Baseline recordings acquired using identical recording parameters enabled direct comparison between baseline and post-stimulation periods. Equivalent baseline recordings were not available for Implant #2 and were therefore excluded from this analysis. GLM analysis demonstrated a significant reduction in IED incidence for all stimulation paradigms compared with baseline (Figure 9D). The greatest reduction was observed during 100 Hz stimulation (IRR = 0.20, 95% CI: 0.13–0.31, p < 0.001), corresponding to an approximately 80% reduction in IED incidence. Stimulation at 50 Hz also produced a substantial reduction (IRR = 0.25, 95% CI: 0.16–0.38, p < 0.001), while 125 Hz stimulation resulted in a more moderate but still significant reduction (IRR = 0.38, 95% CI: 0.25–0.58, p < 0.001).

**Figure 9.**
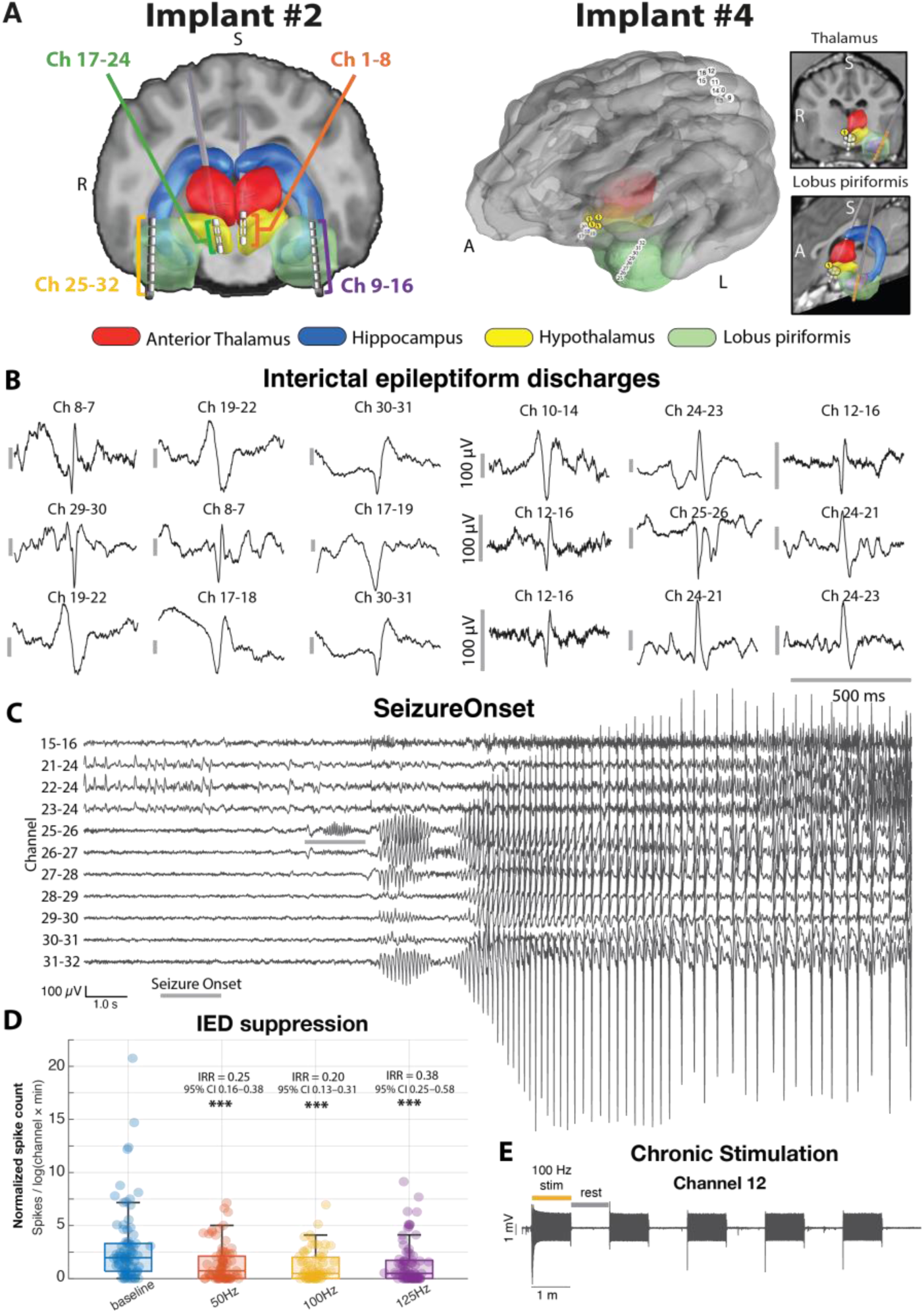
Recording and modulation of epilepsy biomarkers in epileptic canine. **A)** Anatomical reconstructions showing the locations of implanted cortical and depth electrodes in the epileptic canine subjects. **B)** Representative interictal epileptiform discharges (IEDs) recorded from bipolar channel pairs across multiple recording sessions and electrode locations. **C)** Bipolar recordings of a representative seizure captured in Implant #4. The seizure onset zone was localized to channels 25–26 within the piriform lobe, from which ictal activity propagated to other brain regions. **D)** Effect of chronic stimulation on IED incidence in Implant #4. Box plots show normalized IED rates across recording sessions, demonstrating a statistically significant reduction in IED occurrence during stimulation. Statistical analysis was performed using a generalized linear model (GLM). IRR denotes the incidence rate ratio, 95% CI the 95% confidence interval, and asterisks indicate the level of statistical significance (* p < 0.05, ** p < 0.01, *** p < 0.001). **E)** Chronic stimulation protocol. Each stimulation session consisted of a 3-minute baseline recording followed by three 10-minute block stimulation periods delivered at 50, 100, and 125 Hz through the left thalamic depth electrode.

**Brain stimulation evoked potentials** (BSEPs) were successfully recorded following systematic optimization of the recording and stimulation parameters. Contrary to our findings in saline experiments, the best in vivo recordings were obtained without assigning a hardware reference electrode, using only the ground electrode during acquisition and thereby avoiding amplifier saturation during stimulation. Although this recording mode is not formally specified by CorTec, we verified its suitability by acquiring baseline recordings with the same recording configuration in the absence of stimulation. Using these recording parameters, thalamic stimulation elicited reproducible cortical responses, demonstrating effective functional connectivity between the stimulated thalamic nucleus and multiple cortical regions (Figure 10). In the divergent stimulation paradigm, stimulation delivered through the distal contact and the adjacent segmented ring evoked distinct responses in the occipital (Figure 10A), temporal (Figure 10B), and sensorimotor cortices (Figure 10C). In the convergent paradigm, responses were recorded from a single bipolar cortical derivation while varying the stimulation direction within the segmented DBS lead (Figure 10D–G). Changing the active stimulation segments altered both the morphology and amplitude of the evoked responses, with progressively larger responses observed as stimulation was directed toward the recording site.

**Figure 10.**
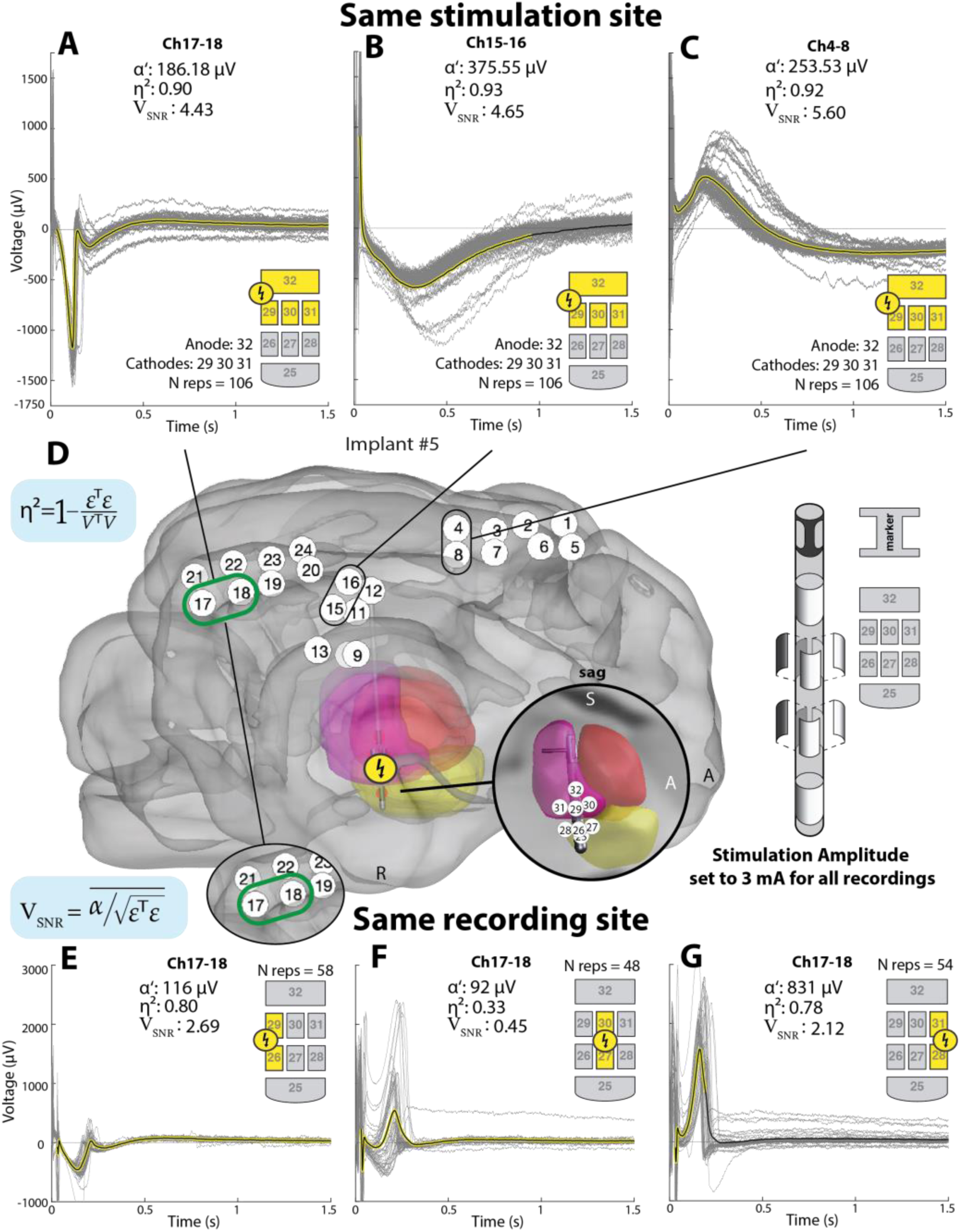
Brain stimulation evoked potentials (BSEPs) recorded with the CorTec Brain Interchange system. A–C) Representative BSEPs recorded from three different recording sites during stimulation delivered through the right thalamic deep brain stimulation (DBS) lead using the distal contact and the adjacent segmented ring. **D) Left:** Subject-specific brain rendering of Implant #5 showing the stimulation and recording electrode locations. **Right:** Schematic of the segmented DBS lead. **E–G)** Representative BSEPs recorded from the same cortical recording site while stimulating in different directions using the three vertical pairs of segmented contacts on the right thalamic DBS lead.

A BCI experiment was performed in a human subject in EMU using the benchtop version of CorTec BIC. Clinical functional motor execution and motor imagery mapping were first performed using the g.tec g.HIamp amplifier to identify cortical sites exhibiting task-related modulation. A bipolar derivation located over the postcentral sulcus demonstrated the strongest imagery-related activity and was subsequently selected for one-dimensional **BCI cursor control** using the benchtop version of the CorTec BIC system (Figure 11A,B).

**Figure 11.**
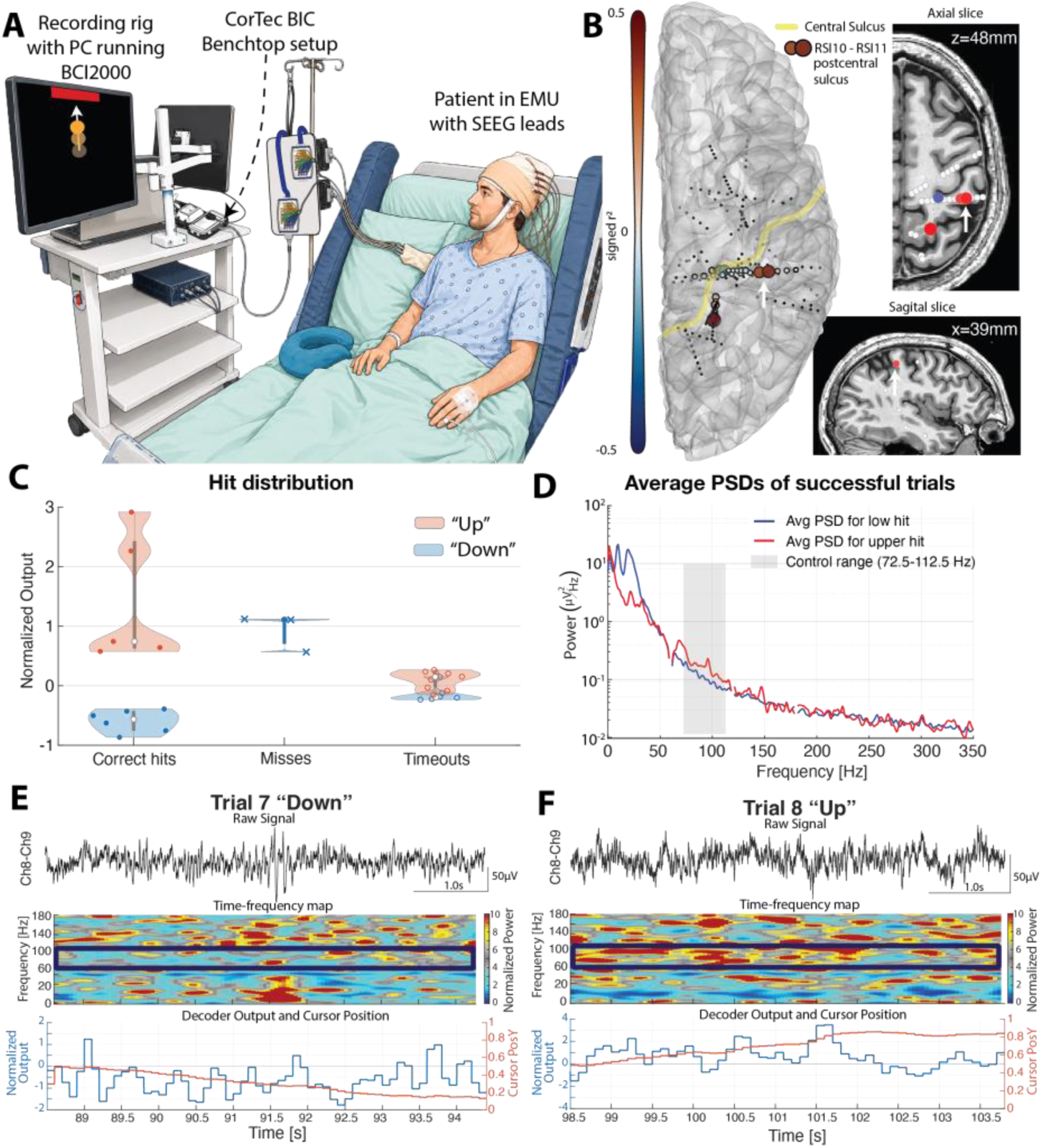
BIC–BCI2000 brain-computer interface (BCI) control using a benchtop Brain Interchange (BIC) device. **A)** Schematic of the recording setup. A patient undergoing stereoencephalography (sEEG) monitoring in the epilepsy monitoring unit (EMU) consented to participate in a research BCI study. The benchtop BIC device was connected to the patient’s externalized sEEG leads while the participant performed a 1D cursor control task. Rectangular targets were presented before cursor movement to cue either movement or rest, and participants attempted to direct the cursor toward the target by modulating their brain activity. The task was performed in the hospital bed while viewing a monitor positioned approximately 80–100 cm from the head. **B)** BCI control was driven by a bipolar electrode pair (indicated by the white arrow) selected based on motor imagery screening performed using a clinical amplifier (g.tec HiAmp) prior to the BCI task. Channel selection was based on the coefficient of determination (r²) calculated from power changes within the 65–115 Hz frequency range. **C)** Normalized output of the online signal processing pipeline for individual cursor control trials, grouped according to trial outcome. During each trial, participants were given 5 seconds to move the cursor either upward or downward by modulating neural activity. Trials in which the target was not reached within the allotted time were classified as Timeout. **D)** Average power spectral densities of successful trials calculated offline. The shaded frequency band (72.5–112.5 Hz) corresponds to the spectral feature used for real-time cursor control feedback during the online BCI task. **E–F)** Representative successful BCI control trials illustrating the raw neural recordings, corresponding time-frequency maps, outputs of the online signal processing pipeline, and the resulting cursor trajectories during real-time control.

The participant completed three overt motor BCI runs followed by one motor imagery run. During the overt BCI runs (30 trials), the participant achieved 11 successful target hits, 3 misses, and 16 timeouts. During the imagery run (26 trials), 5 successful target hits were recorded, with no misses and 21 timeouts (Figure 11C). Representative successful trials are shown in Figure 11E,F. Modulation of high-frequency activity within the control band (72.5–112.5 Hz) was associated with successful cursor movements, while average power spectral densities demonstrated concurrent changes in broadband activity together with modulation of low-frequency power below ∼40 Hz (Figure 11D).

These results demonstrate that an established BCI2000 motor BCI paradigm could be successfully executed using the benchtop CorTec BIC as the signal acquisition device.

## 4. Discussion

We developed and evaluated an open translational ecosystem that integrates a chronically implantable, bidirectional neural interface with the BCI2000 software framework. Three findings are most important. First, quantitative benchtop characterization defines the practical recording, stimulation, and communication performance of the system. Second, longitudinal implantation demonstrates both durable neural recording and identifiable mechanisms of channel degradation.

Third, a common software and hardware framework supported multiple sensing, stimulation, mapping, and BCI paradigms across benchtop, preclinical, and human settings. These findings support the feasibility of the ecosystem for chronic bidirectional neural-interface research and illustrate how shared infrastructure can reduce the effort required to implement new paradigms.

### 4.1. Brain Interchange technical characterization

One of the major goals of this work was to establish quantitative performance metrics that can guide future studies employing the BIC system. While previous publications described the hardware architecture [43], [48], [52] and demonstrated its feasibility for individual applications, including phase-triggered [53] and closed-loop stimulation [84], bedside detection of IEDs and HFOs [48] [85], cortico-cortical [84] and somatosensory evoked potentials [54], comprehensive characterization of recording performance, transfer characteristics, impedance measurements, and intrinsic communication latency remained limited. The present work addresses several of these gaps and provides a reference dataset for future investigations.

The benchtop characterization demonstrated that higher amplifier gains progressively reduced the recording noise and are therefore preferable for passive neural recordings, whereas lower gains reduce amplifier saturation during stimulation experiments. Although the effective recording bandwidth (approximately 2–325 Hz) is sufficient for most intracranial biomarkers, including EEG rhythms, epileptiform spikes, and broadband activity, applications targeting high-frequency oscillations (HFOs) or fast ripples may require additional signal processing techniques [79], [86].

Similarly, impedance measurements provide reliable estimates within the physiologically relevant range encountered during chronic recordings and should primarily be interpreted as indicators of electrode integrity rather than precise electrical measurements. We further demonstrate that the choice of hardware and software referencing strategy depends on the intended application and can substantially influence the recorded signals. Bipolar re-referencing reduces the amplitude of coherent oscillatory activity while increasing the broadband noise floor. In contrast, common average reference (CAR) methods reduce the broadband noise floor but also alter the amplitude of coherent oscillatory components. A detailed evaluation of these effects is provided in the supplement, and practical recommendations for selecting appropriate referencing strategies are presented in our previous work [87].

Another important contribution is the characterization of the intrinsic acquisition and stimulation latency of the BIC system. Unlike previous reports [50], [51], [53], [88], which quantified the combined overall system latency, we separately characterized the acquisition and stimulation latencies, establishing a theoretical hardware lower bound of approximately 22 ms for closed-loop interventions. This provides a reference baseline against which future closed-loop implementations can be evaluated. In practice, however, the overall response time will depend on the selected signal processing pipeline, requiring investigators to balance detection complexity against response latency.

The present study also provides the longest longitudinal evaluations of the BIC platform to date, with five large animal implants monitored in total for more than three years and the longest follow-up exceeding 1,000 days. Compared with the approximately 400-day BIC chronic recordings reported by Yang et al. [54], this substantially extends the duration of long-term validation while also demonstrating, for the first time, chronic use of both ECoG and DBS electrodes with the BIC system.

Throughout the implantation period, wireless communication remained stable, supporting the feasibility of long-term recordings. Continuous monitoring also provided valuable insight into the mechanisms underlying channel degradation while highlighting the challenges of assessing channel quality during gradual long-term deterioration. Unlike abrupt hardware failures, slowly degrading channels often transition over extended periods between clearly physiological and clearly artifactual recordings, making it difficult to define the exact time point at which a channel becomes unusable. We addressed this challenge by empirically and algorithmically assessing signal usability through a combination of impedance measurements, temporal and spectral signal characteristics, and follow-up CT imaging. This approach enabled identification of several mechanical causes of recording failure, including electrode pull-out, strip folding, connector misalignment, lead migration, and wire breakage. Beyond supporting long-term monitoring, these observations inform future studies by highlighting common failure mechanisms and practical considerations for chronic implantation and data quality assessment.

### 4.2. General-purpose translational ecosystem

Perhaps the most important outcome of this work is the demonstration that a single hardware and software ecosystem can support a broad spectrum of neuromodulation paradigms without requiring substantial modifications to the underlying infrastructure. Rather than representing isolated applications, the presented results collectively demonstrate the versatility of the ecosystem and its ability to support the complete translational workflow, from biomarker discovery and functional localization to stimulation optimization and human proof-of-concept validation. The ecosystem was designed so that individual hardware and software components can be readily exchanged or extended to meet the requirements of different research applications. The works of Dold et al. [50] and Afshar et al. [89] are conceptually the closest to our approach, as they similarly promote open, device-agnostic software frameworks for neuromodulation research. Our work, however, builds upon the mature BCI2000 platform, leveraging its validated signal-processing modules, synchronization tools, visualization utilities, and established user community.

Using the same platform, we demonstrated functional mapping, chronic neural sensing and biomarker monitoring, adaptive and open-loop stimulation, BSEPs, and implantable BCI control. Functional recordings continued to capture physiologically meaningful cortical activity that closely corresponded with established canine neuroanatomy even after more than two years post implantation. Chronic recordings in a canine with naturally occurring epilepsy enabled capture of seizure, localization of the SOZ, longitudinal monitoring of IEDs, and evaluation of their modulation by thalamic stimulation. The same ecosystem also enabled recording of BSEPs, with directional DBS producing distinct evoked responses. Furthermore, the modular architecture enabled implementation of closed-loop stimulation based on arbitrary user-defined neural features, while providing a framework for balancing detection accuracy and response latency according to the requirements of a given application. Lastly, we demonstrate the practical portability of the ecosystem by translating an established BCI2000 motor BCI paradigm to the Brain Interchange platform with only minimal modifications to the existing signal-processing pipeline.

This versatility is particularly important because future adaptive neuromodulation research increasingly requires investigators to move seamlessly between biomarker identification, network mapping, optimization of stimulation strategies, and clinical validation. Rather than developing dedicated hardware and software solutions for each individual application, the presented ecosystem provides a common experimental infrastructure that supports all of these paradigms within a unified framework. Together, these demonstrations show that sensing, stimulation, biomarker evaluation, and neural feedback control should not be viewed as independent research directions, but as complementary components of a single translational ecosystem capable of accelerating the iterative development of next-generation neuromodulation therapies.

### 4.3. Consideration of current and future directions

Several limitations of the present study should be acknowledged while also highlighting opportunities for future development. The preclinical validation was performed in a limited number of animals, including only a single canine with naturally occurring epilepsy, and the human evaluation consisted of a single proof-of-concept participant. Consequently, several demonstrated paradigms, including chronic biomarker modulation and implantable BCI control, should be regarded as demonstrations of feasibility rather than clinical efficacy. Validation in larger preclinical and clinical cohorts will be required to establish their safety, therapeutic efficacy, robustness, and generalizability.

The canine model also introduced practical challenges that differ from the intended clinical application, including greater mechanical stress on implanted leads, smaller neuroanatomical targets, intermittent detachment of the magnetically coupled headpiece, and the need for inductive powering, which limited recordings to experimental sessions rather than continuous monitoring. While these factors complicate long-term evaluation, they primarily reflect characteristics of the preclinical model rather than inherent limitations of the underlying technology. The mechanical failure mechanisms identified in this study informed subsequent hardware refinements, leading to the adoption of more mechanically robust stranded lead wires in newer electrode generations.

The proof-of-concept closed-loop stimulation paradigm demonstrated relatively long response latencies because the selected algorithm relied on spectral estimation over long temporal windows. While this implementation prioritized robustness against noise and stimulation artifacts, alternative signal processing approaches based on time-domain features can substantially reduce latency when appropriate for the intended application. Likewise, the stimulation-evoked potentials presented here were acquired using a recording configuration optimized to minimize amplifier saturation during stimulation. Reliable acquisition of BSEPs remains an ongoing challenge, highlighting the need for continued refinement of the BIC hardware, stimulation protocols, and signal processing to improve artifact suppression, recording robustness, latency, and signal quality.

Similarly, the demonstrated implantable BCI paradigm achieved only modest decoding performance because the participant had limited training time and electrode placement was dictated entirely by clinical considerations rather than BCI optimization. Future studies employing electrodes specifically positioned for motor decoding, together with personalized decoding algorithms and extended user training, are expected to substantially improve performance.

More broadly, the open architecture presented here provides a foundation for continued development of adaptive neuromodulation applications, including on-the-fly spike detection, responsive neurostimulation, distributed adaptive DBS, and increasingly sophisticated implantable BCIs. As these algorithms mature, computational components currently implemented within BCI2000 can progressively be optimized and migrated to embedded implementations while maintaining compatibility with the surrounding research ecosystem. In parallel, we are collaborating with CorTec on a next-generation miniaturized external unit that integrates the communication unit and headpiece into a single compact device. Besides improving usability, reducing the distance between the inductive power and communication antennas is expected to improve wireless reliability and facilitate longer recordings in natural environments using portable edge-computing devices (Supplementary Figure 16).

## 5. Conclusion

We present an open, modular ecosystem integrating the CorTec Brain Interchange with BCI2000 for implantable BCI and adaptive neuromodulation research. Benchtop, chronic preclinical, and human proof-of-concept studies demonstrate that the framework can support neural recording, functional mapping, biomarker detection, stimulation, evoked-potential recording, and BCI control. The accompanying technical characterization, software, datasets, and experimental workflows provide a reproducible foundation for future development and evaluation of implantable neurotechnology. Ultimately, we hope these efforts will accelerate the development of next-generation implantable neuromodulation technologies and support personalized therapies for psychiatric and neurological disorders such as epilepsy, Parkinson’s disease, and amyotrophic lateral sclerosis.

## Acknowledgements

This work was made possible with the support of the National Institutes of Health (NIH), whose funding is gratefully acknowledged, specifically through the BRAIN Initiative award U01NS128612 (KJM, GAW, PB). Additional support was provided by the NIH awards and grants NCATS CTSA KL2-TR002379 (KJM), NIBIB P41-EB018783 & R01-EB026439 (PB), U24-NS109103 (PB), UH3NS117944 (NFI), R01NS112497 (NFI), UH2&3 NS095495 (GAW), and R01-NS92882 (GAW) from the BRAIN Initiative and National Institute of Neurological Disorders and Stroke (NINDS). DJK was additionally supported by the Mayo Clinic T32 training grant (T32 GM145408). The contents of this manuscript are solely the responsibility of the authors and do not necessarily represent the official views of the NIH. Our funders played no role in data collection and analysis, study design, decision to publish, or manuscript preparation.

## Manuscript preparation

This is the version of the article before peer review or editing, as submitted by an author to IOP Journal of Neural Engineering. IOP Publishing Ltd is not responsible for any errors or omissions in this version of the manuscript or any version derived from it. Parts of this manuscript were revised and edited with the assistance of OpenAI’s ChatGPT, a large language model, to improve clarity and language. Elements of figures [1, 4, 7, 11] were partly generated using ChatGPT’s image generation capabilities. All intellectual content, scientific conclusions, and final wording were determined and approved by the authors.

## Data and code availability statement

The data that support the findings of this study are openly available at https://openneuro.org/datasets/ds004624/versions/2.0.0 & https://dandiarchive.org/dandiset/000571/0.250616.1143?search=BCI2000&pos=1. All code is available at the GitHub repostiroty: https://github.com/Cybernetics-and-Motor-Physiology-Lab/BIC-BCI2000_ecosystem. A Comprehensive documentation, tutorials, software modules, example datasets, and surgical protocols are freely available through the BCI2000 Wiki (https://www.bci2000.org/mediawiki/index.php/CortecExperience) & https://www.bci2000.org/mediawiki/index.php/Contributions:CortecADC.

## Author contributions

F.L., P.B., G.A.W., G.S., and K.J.M.. designed research; F.L., F.M., A.H.A., M.vd.B., J.B., D.J.K., and K.J.M. performed research and collected the data; F.L., F.M., N.L., W.E., A.H.A., B.F.B., I.K.,N.F.I., P.B., and K.J.M. contributed new reagents/analytic tools; F.L., M.vd.B., W.E. and K.J.M. analyzed data; F.L., M.R.B., and K.J.M. wrote the original draft; and V.K., N.P.S., G.S., N.F.I., P.B., AND G.A.W. provided review and guidance in research and the editing of the paper.

## Footnotes

1 In compliance with the Minnesota state Beagle Freedom Bill statute 135A.191.

