## Supplemental Data 1 for "A Translational Platform for Brain-Computer Interfaces and Adaptive Neuromodulation: Technical Characterization, Long-Term Validation, and Implementation of the CorTec Brain Interchange–BCI2000 Ecosystem"

### Supplement

#### 1. Abbreviations

| Abbreviation | Definition |
|---|---|
| 95% CI | 95% confidence intervals |
| ANT | Anterior nucleus of the thalamus |
| API | Application Programming Interface |
| BIC | CorTec Brain Interchange |
| BCI | Brain-computer interface |
| BP | Bi-polar rereferencing |
| BSEPs | Brain stimulation evoked potentials |
| CAR | common average reference |
| CT | Computed tomography |
| DBS | deep brain stimulation |
| ECoG | electrocorticography |
| EMU | Epilepsy monitoring unit |
| GLM | generalized linear model |
| HFOs | High-frequency oscillations |
| HPC | Hippocampus |
| IEDs | Interictal epileptiform discharges |
| IRR | Incidence rate ratios |
| LeadCAR | lead common average rereferncing |
| MRI | Magnetic resonance imaging |
| PL | Packet loss |
| PSD | Power spectral density |
| RMS | Root mean square value |
| sEEG | Stereoelectroencephalography |
| SOZ | Seizure onset zone |
| SPES | Single-pulse electrical stimulation |
| TTL | Transistor-Transistor Logic |

#### 2. In-vitro testing

##### Frequency sweep

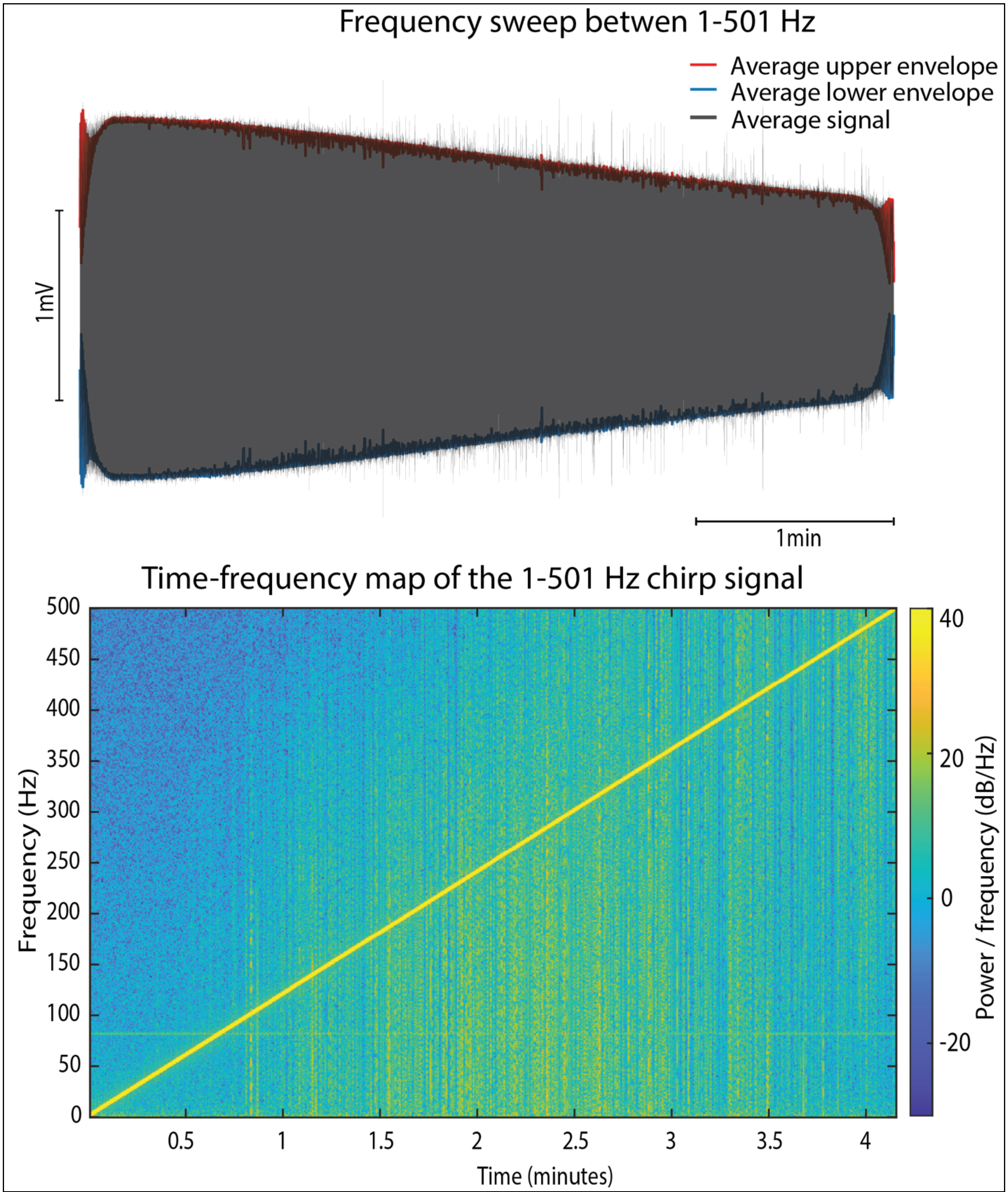

**Figure 1. Frequency sweep protocol.** The amplifier transfer function was evaluated using a 2 mV peak-to-peak chirp signal sweeping from 1 to 501 Hz over 250 s. **Top**: Input chirp waveform and its amplitude envelope. **Bottom**: Time-frequency representation of the input signal.

#### BIC Latency Evaluation

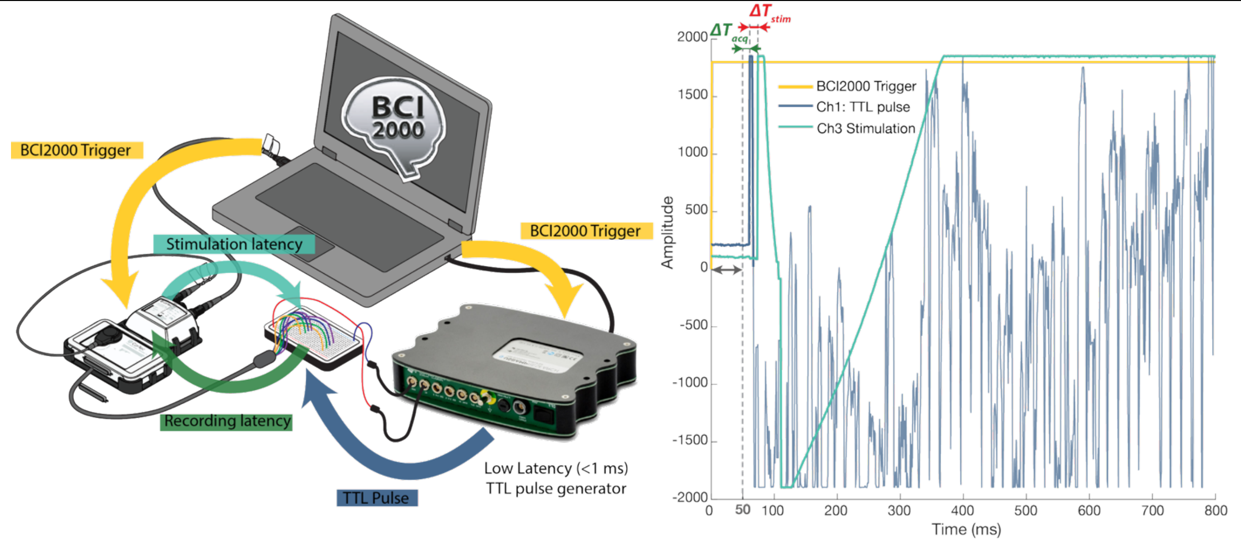

**Figure 2. Latency evaluation. Left:** Schematic of the latency measurement setup. BCI2000 simultaneously sends a stimulation trigger to the CorTec BrainInterchange (BIC) system and to a low-latency (<1 ms) TTL pulse generator. The TTL pulse is recorded by the BIC, providing a ground-truth reference for acquisition timing, while the BIC delivers a stimulation pulse in response to the same trigger. **Right**: Representative recording illustrating latency measurements. Acquisition latency ($\Delta T_{acq}$) was defined as the interval between the BCI2000 trigger and the recorded TTL pulse. Stimulation latency ($\Delta T_{stim}$) was defined as the interval between the BCI2000 trigger and the detected stimulation artifact.

#### Impedance measurement pulses

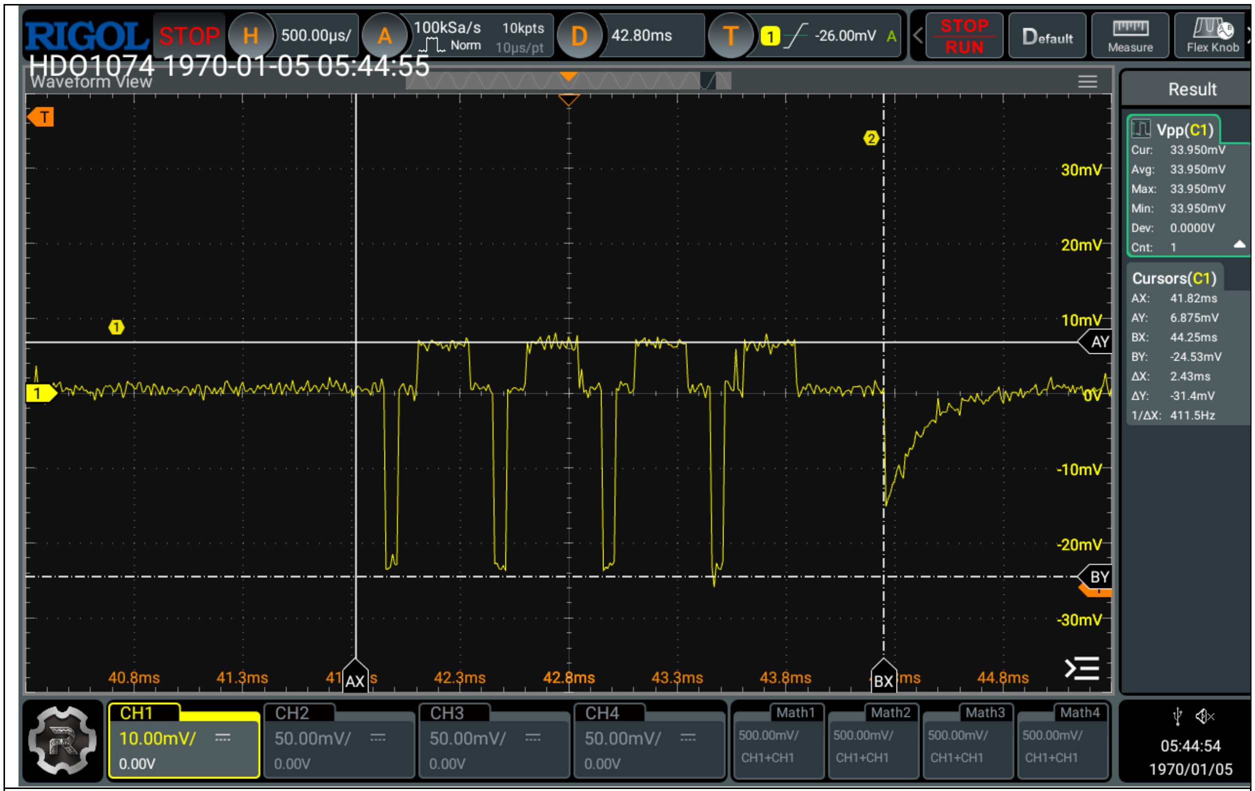

**Figure 3. Impedance measurement pulses**. Representative impedance measurement pulse recorded with a laboratory oscilloscope across a 1 kΩ precision resistor.

#### Impedance measurement results

| Connector impedance [Ω] | Wire impedance [Ω] |
|---|---|
| 0.25+0.21 | 0.025 |

| Resistors | | | | | |
|---|---|---|---|---|---|
| Declared value [Ω] | 100 | 470 | 1000 | 2200 | 560 k |
| Actual Value [Ω] | 99.6 | 471.5 | 997.7 | 2152 | 560.7 k |

| Capacitors | | | |
|---|---|---|---|
| Declared value [nF] | 150 | 220 | 330 |
| Actual Value [nF] | 140 | 218 | 324.5 |

| Pure resistance | |
|---|---|
| Inserted R [Ω] | 997.7 |

| | |
|---|---|
| | 1040 |
| | 1015 |
| | 1011 |
| | 969 |
| | 1036 |
| | 1022 |
| | 1038 |
| | 1011 |
| | 1020 |
| | 1029 |
| Mean | 1019.1 |
| STD | 19.61 |

| Varying resistance (C set to 220nF) | | | | |
|---|---|---|---|---|
| Inserted R [Ω] | 100 | 470 | 1k | 2k |
| Calculated \|Z\| [Ω] | 100 | 472 | 998 | 2152 |
| | 23 | 412 | 870 | 2047 |
| | 46 | 385 | 877 | 2020 |
| | 55 | 408 | 921 | 2071 |
| | 36 | 380 | 916 | 2091 |
| | 50 | 420 | 868 | 2026 |
| | 71 | 404 | 952 | 2033 |
| | 17 | 389 | 904 | 2060 |
| | 55 | 403 | 910 | 2075 |
| | 46 | 439 | 918 | 2081 |
| | 23 | 393 | 956 | 2092 |
| Mean | 42.2 | 403.3 | 909.2 | 2059.6 |
| STD | 16.30 | 16.85 | 29.30 | 25.35 |

| Changing the capacitance (R set to 470Ω) | | | |
|---|---|---|---|
| Inserted C | 150 | 220 | 330 |
| Calculated \|Z\| [Ω] | 472 | 472 | 472 |
| | 300 | 412 | 418 |
| | 324 | 382 | 418 |
| | 326 | 406 | 427 |
| | 322 | 380 | 423 |
| | 376 | 427 | 422 |

| | | | |
|---|---|---|---|
| | 343 | 404 | 444 |
| | 340 | 389 | 452 |
| | 347 | 403 | 429 |
| | 351 | 439 | 450 |
| | 319 | 393 | 464 |
| Mean | 334.8 | 403.5 | 434.7 |
| STD | 20.05 | 17.96 | 15.58 |

#### Stimulation Pulse definition

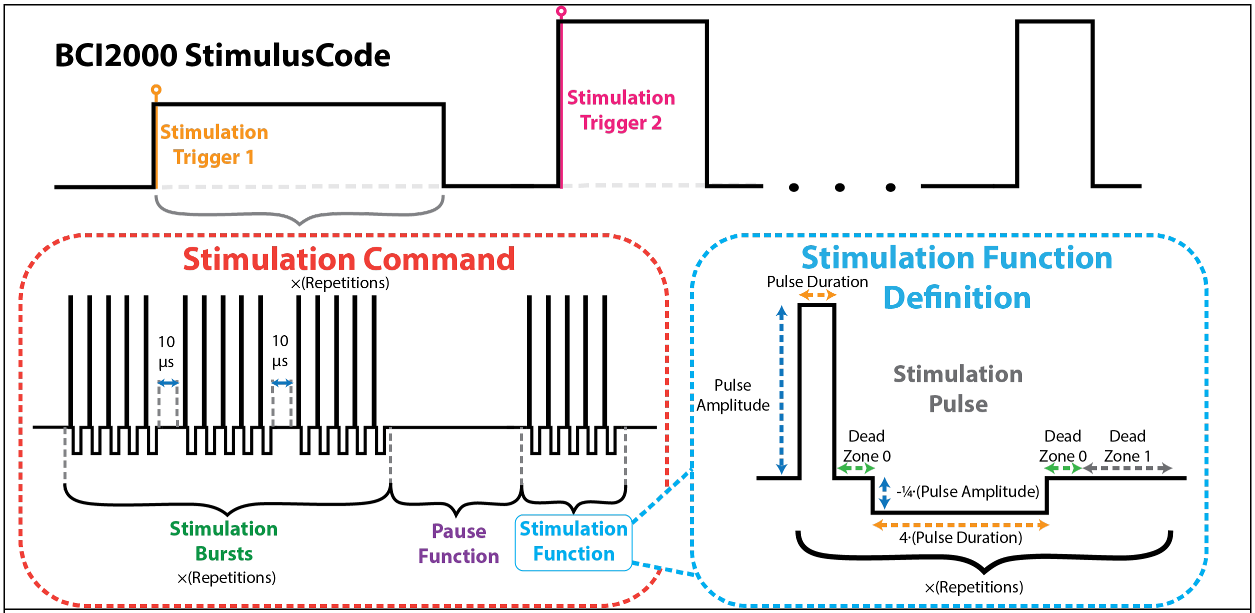

**Figure 4. CorTec Brain Interchange (BIC) stimulation pulse definition in BCI2000**. A stimulation command is initiated by changes in the BCI2000 StimulusCode state variable and consists of one or more stimulation or pause functions. Each stimulation function is composed of repeated biphasic asymmetric pulses, with user-defined pulse amplitude, pulse duration, interphase dead zone, and repetition count.

#### Stimulation Preloading Modes

##### *Stimulation Volatile Command Preloading mode*

As the name suggest, in this stimulation preloading mode the stimulation command can be set arbitrarily big. The stimulation command will be executed in order starting with the first stimulation function and ending with the final stimulation function. Whenever a stimulation function has been executed, it will be then removed from the queue. This implies that an enqueued stimulation command will be executed exactly just one time.

##### *Stimulation Persistent Command Preloading mode*

On the contrary to the volatile command preloading mode, the stimulation command can not be picked arbitrarily large. The stimulation command on this mode has the constraint, that the number of the enqueued stimulation functions must not exceed 16. Yet, in the same manner as the volatile command preloading mode the stimulation command will be executed in order starting from the first stimulation function. The stimulation functions however will not be removed after their execution. Consequently, starting the same stimulation command multiple times in a row will lead to no problems.

##### *Stimulation Persistent Function preloading mode*

In this stimulation preloading mode, the stimulation command can not be picked arbitrarily large. The stimulation command on this mode has the constraint, that the number of the enqueued stimulation functions must not exceed 16. The stimulation functions similar to the stimulation command preloading mode case will not be removed after their execution. However, only one of the stimulation command's function can be executed at one time. Consequently, starting the same stimulation function multiple times in a row will lead to no problems. We can select the stimulation function to be executed by calling start stimulation with the stimulation function index. Remark : The Stimulation command repetitions in this mode will be ignored. Only the selected stimulation function will be executed.

#### Stability testing

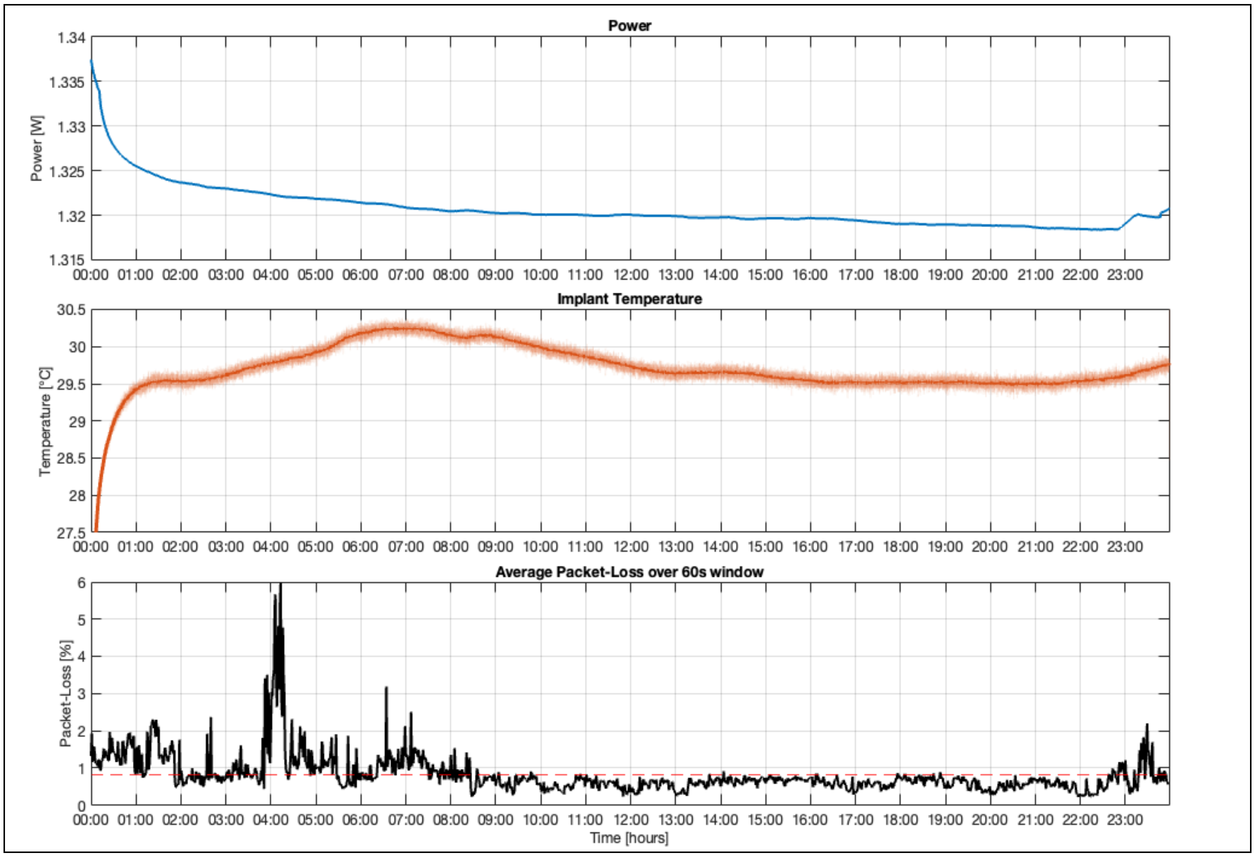

**Figure 5. BIC recording stability**. Long-term stability of the bench-top Brain Interchange (BIC) system during continuous 24-hour operation under laboratory conditions. From top to bottom: device power consumption, implant temperature, and average wireless packet loss calculated over 60-second windows. Power consumption was monitored using an FNIRSI USB power logger.

#### Hardware and Software Re-referencing

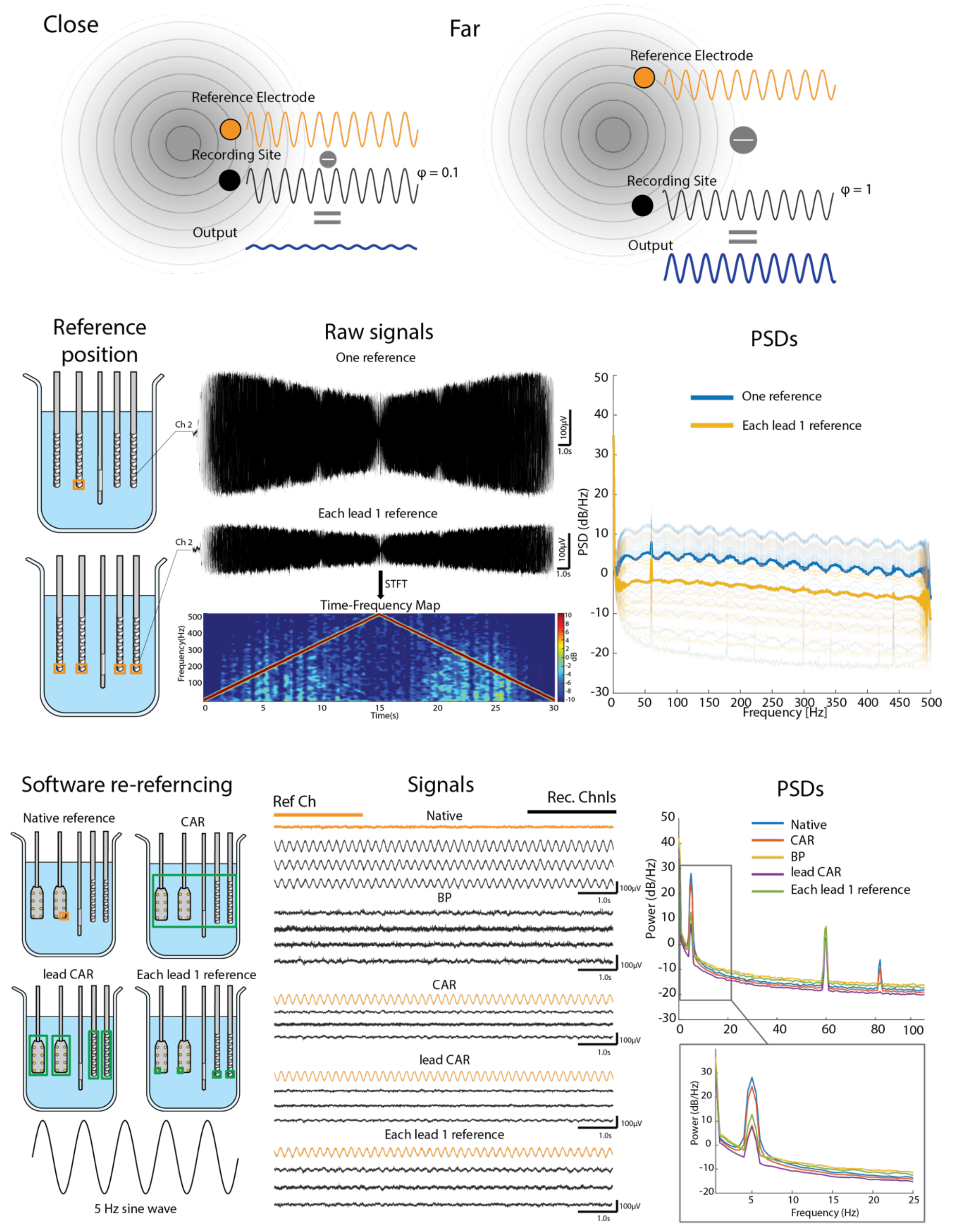

**Figure 6.** Effects of hardware and software referencing strategies. **Top:** Schematic illustrating the principle of common-mode rejection. Signals with similar phase recorded at the recording and reference electrodes

are attenuated by differential (bipolar) recording, whereas signals with larger spatial phase differences are preserved. This demonstrates the effectiveness of bipolar referencing for suppressing spatially coherent interference while retaining local neural activity. **Middle:** Influence of hardware reference electrode selection on signal amplitude and power spectral density, demonstrating the effect of common-mode rejection. **Bottom:** Representative example of the effects of different software rereferencing strategies on a 5 Hz sinusoidal signal, shown in the time and frequency domains.

#### 3. In-vivo testing

##### Image co-registration

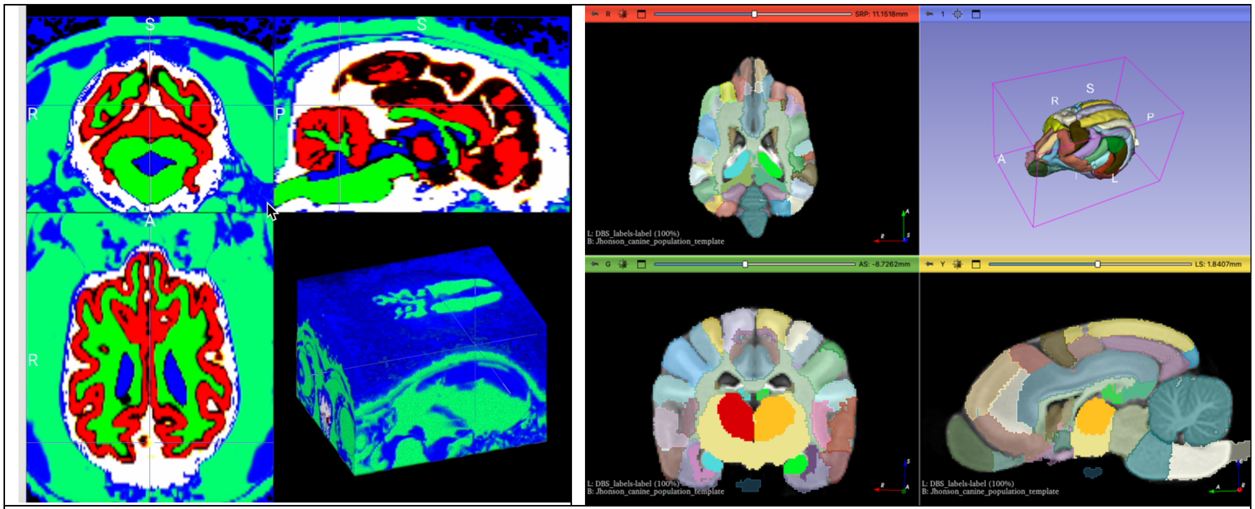

**Figure 7. Canine imaging pipeline**. **Left:** Tissue segmentation of canine brain MRI using tissue probability maps (TPMs) implemented in SPM. **Right:** Custom canine deep brain stimulation (DBS) atlas created by integrating anatomical labels from the Johnson and Czeibert canine brain atlases into a unified template.

#### Implant #1 long-term recording capabilities

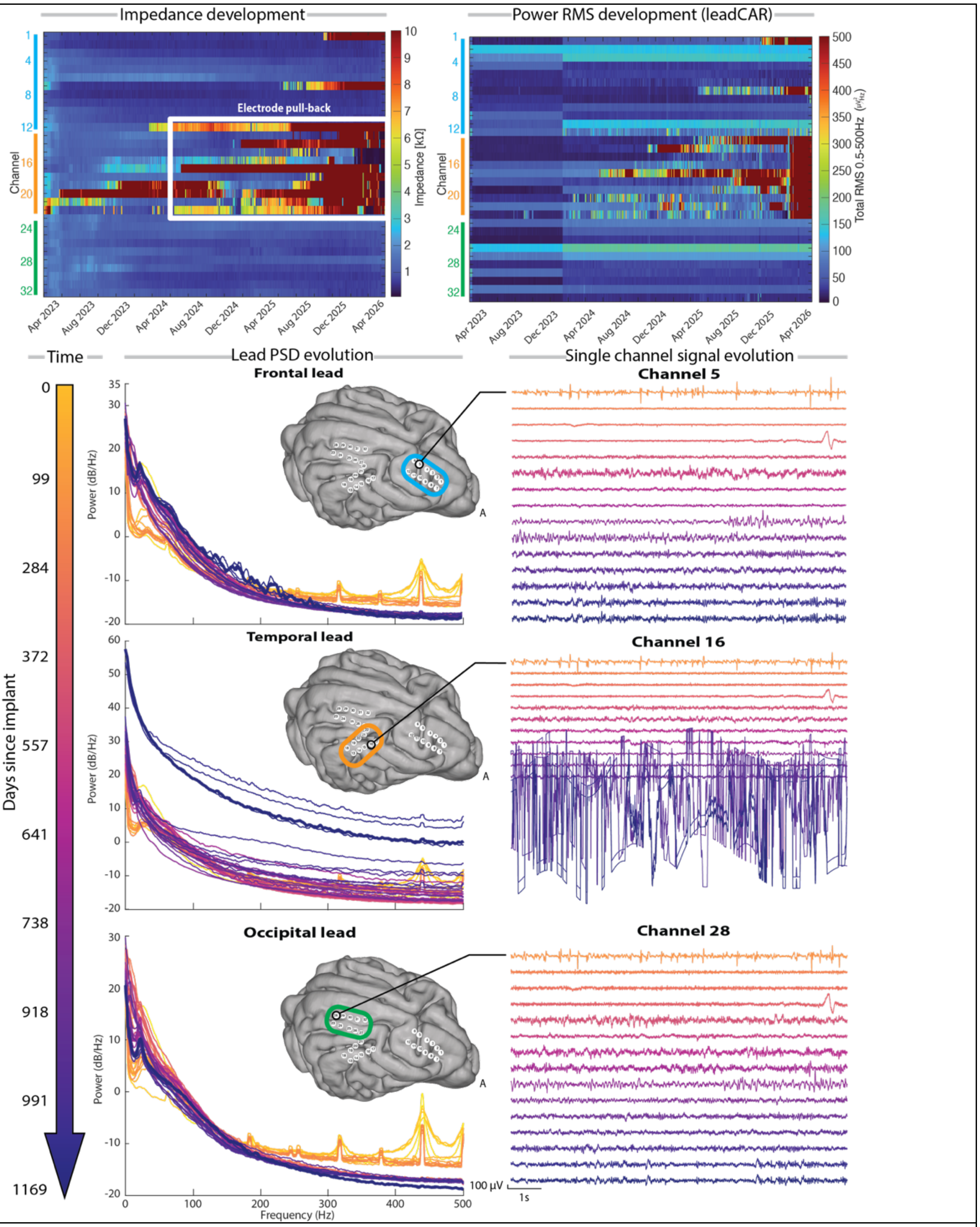

**Figure 8**. Long-term recording performance of Implant #1. **Top left**: Longitudinal electrode impedance measurements across all recording channels. The highlighted region indicates a period of mechanical electrode pull-back. **Top right**: Root mean square (RMS) signal amplitude derived from passive recordings after lead common average rereferencing (leadCAR). **Bottom**: Longitudinal evolution of the median power spectral density (PSD) for each implanted lead, calculated from passive recordings using the native hardware reference. Representative signal traces from one recording channel within each lead illustrate stable (Channels 5 and 28) and degraded (Channel 16) recording performance over the implantation period.

#### Implant #2 long-term recording capabilities

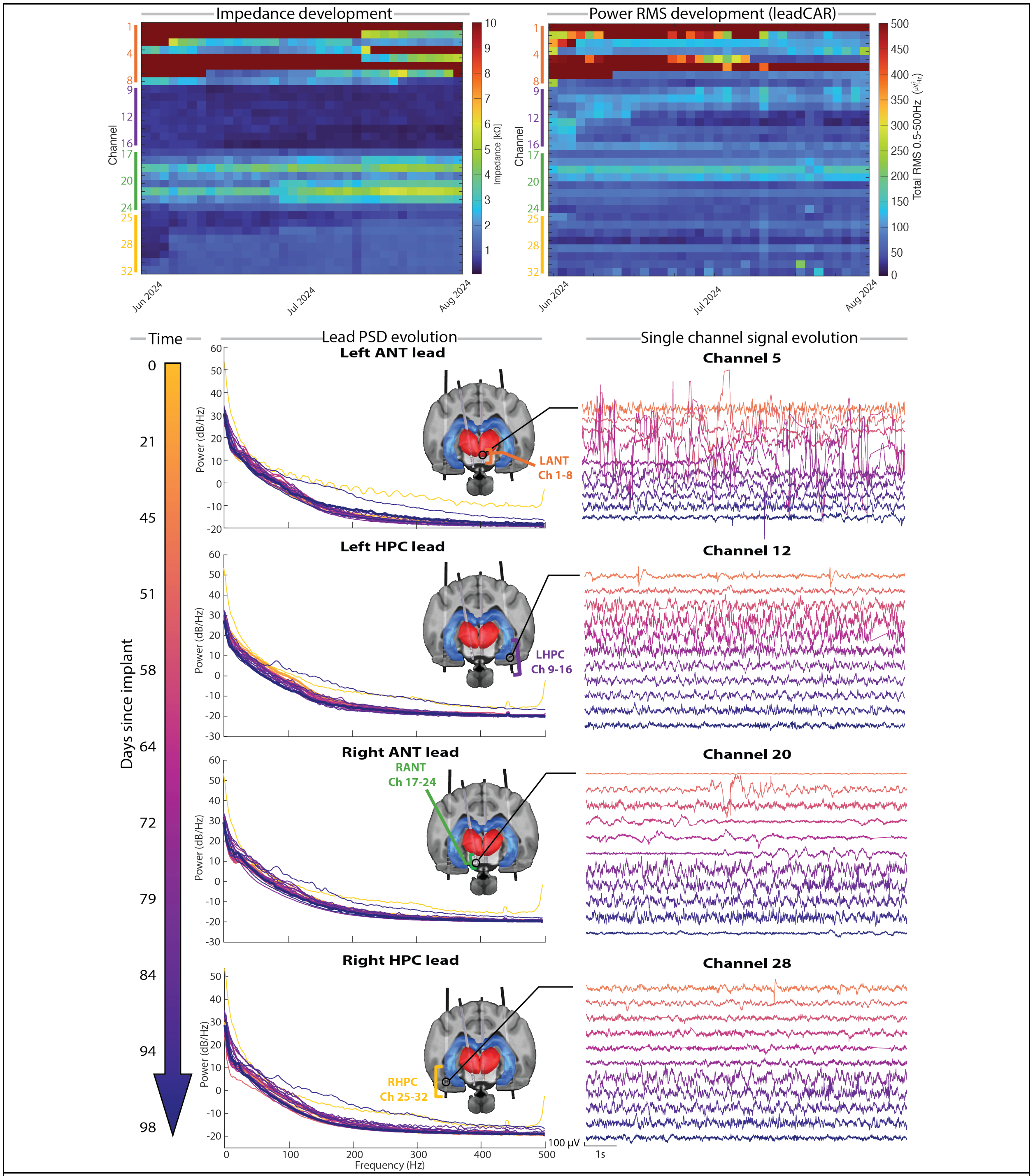

**Figure 9**. **Long-term recording performance of Implant #2. Top left**: Longitudinal electrode impedance measurements across all recording channels. **Top right**: Root mean square (RMS) signal amplitude derived from passive recordings after lead common average rereferencing (leadCAR). **Bottom**: Longitudinal evolution of the median power spectral density (PSD) for each implanted lead, calculated from passive recordings using the native hardware reference. Representative signal traces from one recording channel within each lead illustrate the stability of the recorded neural signals throughout the implantation period.

#### Implant #3 long-term recording capabilities

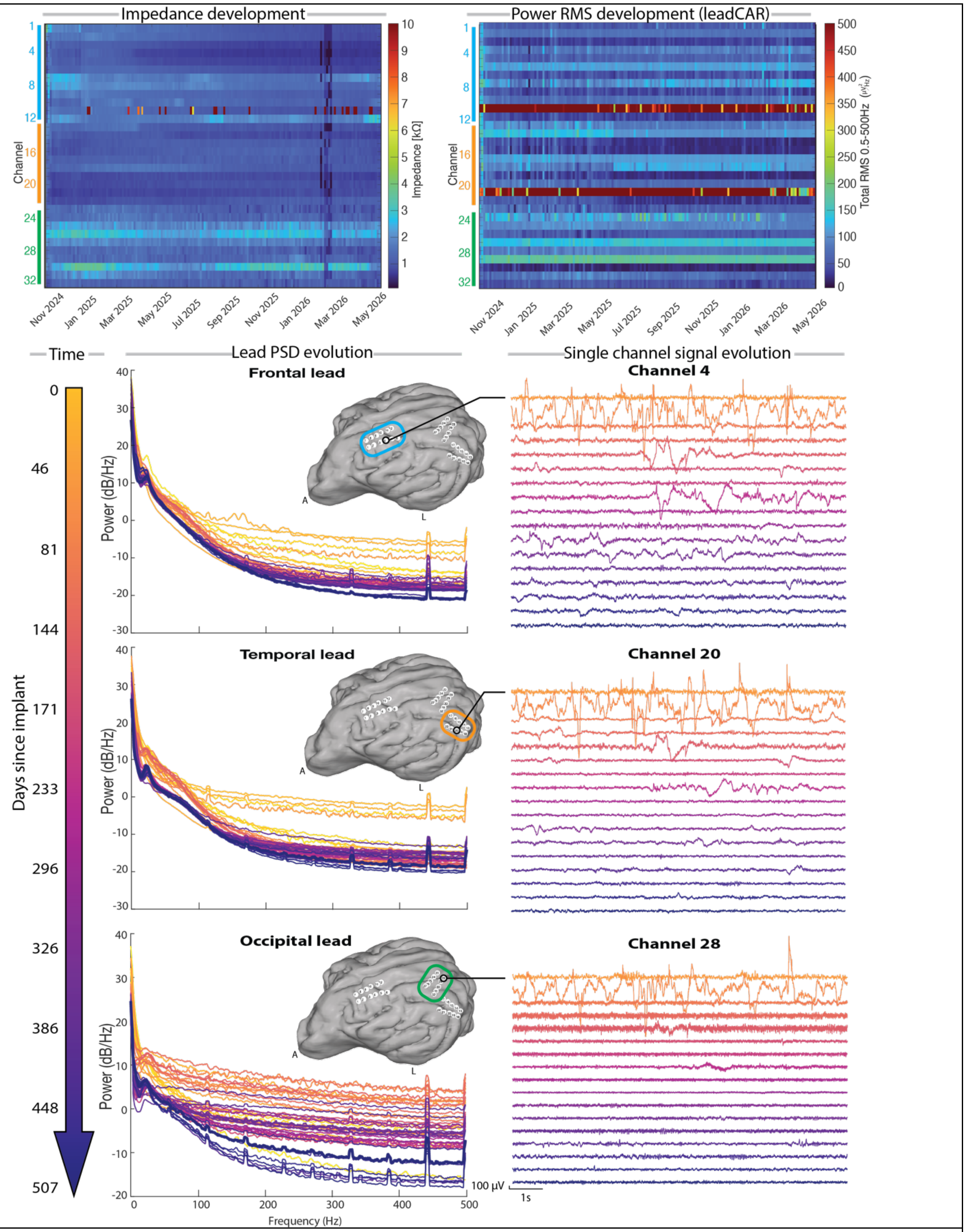

**Figure 10. Long-term recording performance of Implant #3**. **Top left**: Longitudinal electrode impedance measurements across all recording channels. **Top right**: Root mean square (RMS) signal amplitude derived from passive recordings after lead common average rereferencing (leadCAR). **Bottom**: Longitudinal evolution of the median power spectral density (PSD) for each implanted lead, calculated from passive recordings using the native hardware reference. Representative signal traces from one recording channel within each lead illustrate the stability of the recorded neural signals throughout the implantation period.

#### Implant #4 long-term recording capabilities

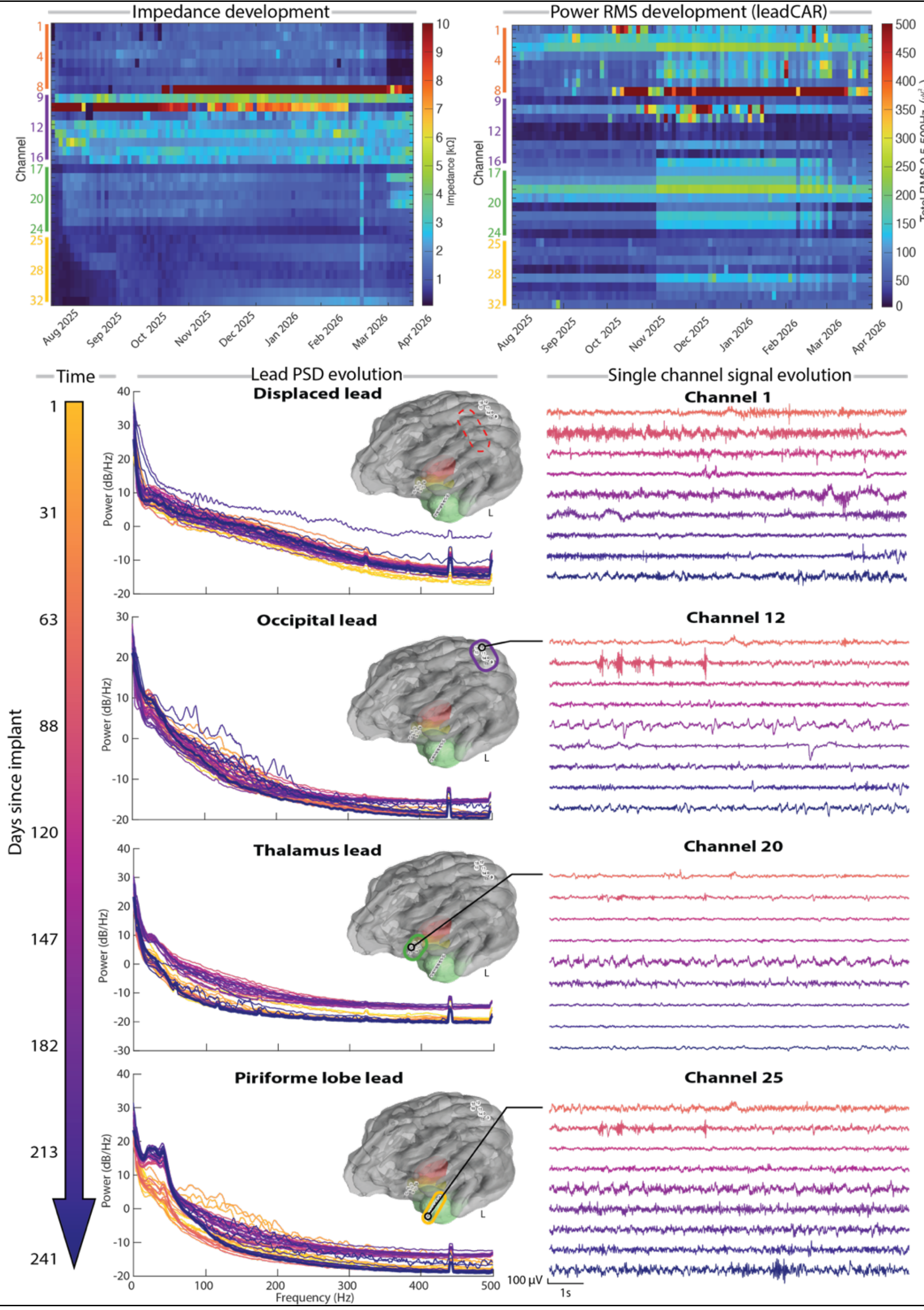

**Figure 11. Long-term recording performance of Implant #4. Top left**: Longitudinal electrode impedance measurements across all recording channels. **Top right**: Root mean square (RMS) signal amplitude derived from passive recordings after lead common average rereferencing (leadCAR). **Bottom**: Longitudinal evolution of the median power spectral density (PSD) for each implanted lead, calculated from passive recordings using the native hardware reference. Representative signal traces from one recording channel within each lead illustrate the stability of the recorded neural signals throughout the implantation period.

#### Implant #5 long-term recording capabilities

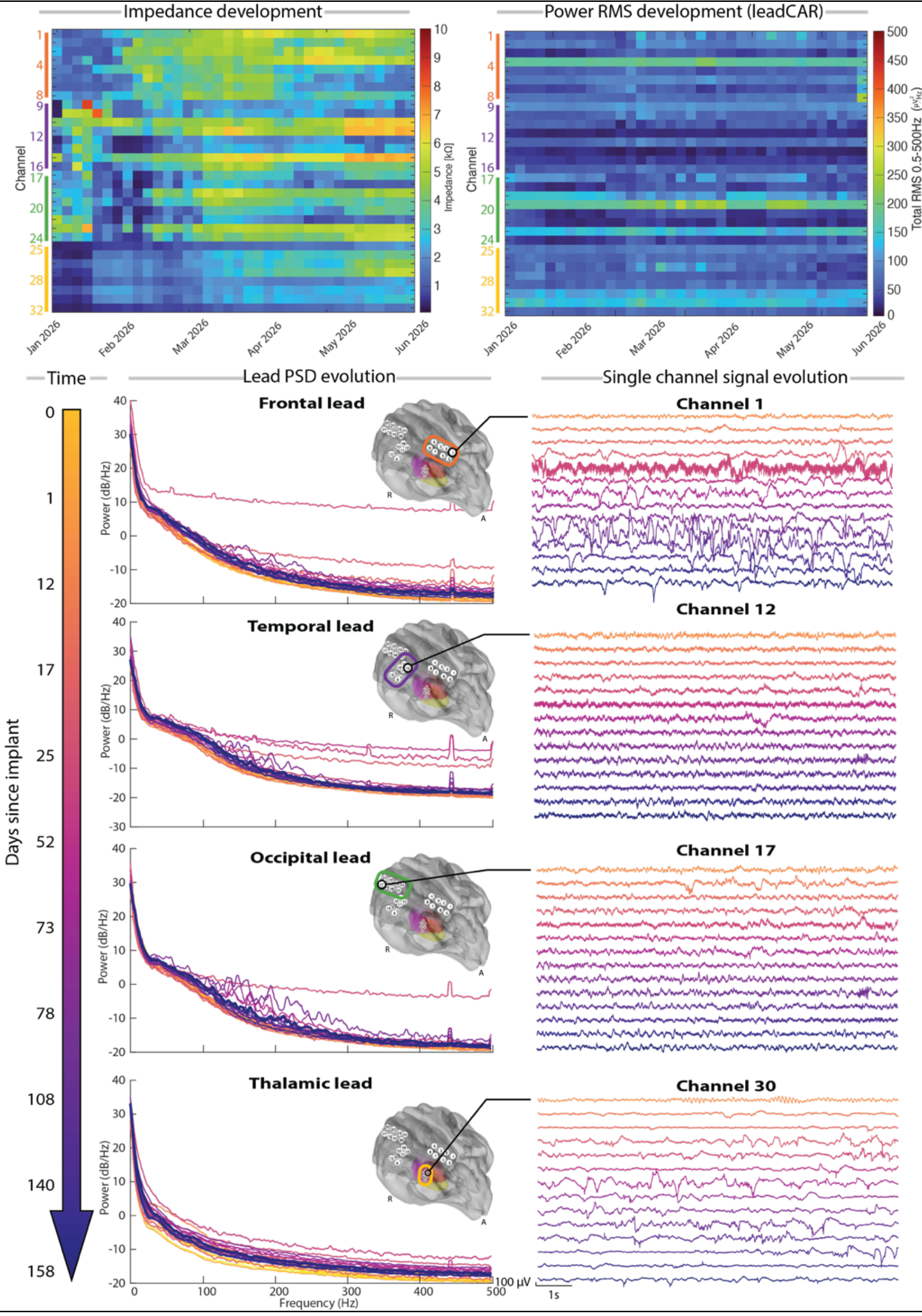

**Figure 12. Long-term recording performance of Implant #5**. **Top left**: Longitudinal electrode impedance measurements across all recording channels. **Top right**: Root mean square (RMS) signal amplitude derived from passive recordings after lead common average rereferencing (leadCAR). **Bottom**: Longitudinal evolution of the median power spectral density (PSD) for each implanted lead, calculated from passive recordings using the native hardware reference. Representative signal traces from one recording channel within each lead illustrate the stability of the recorded neural signals throughout the implantation period.

#### Bad channel detection

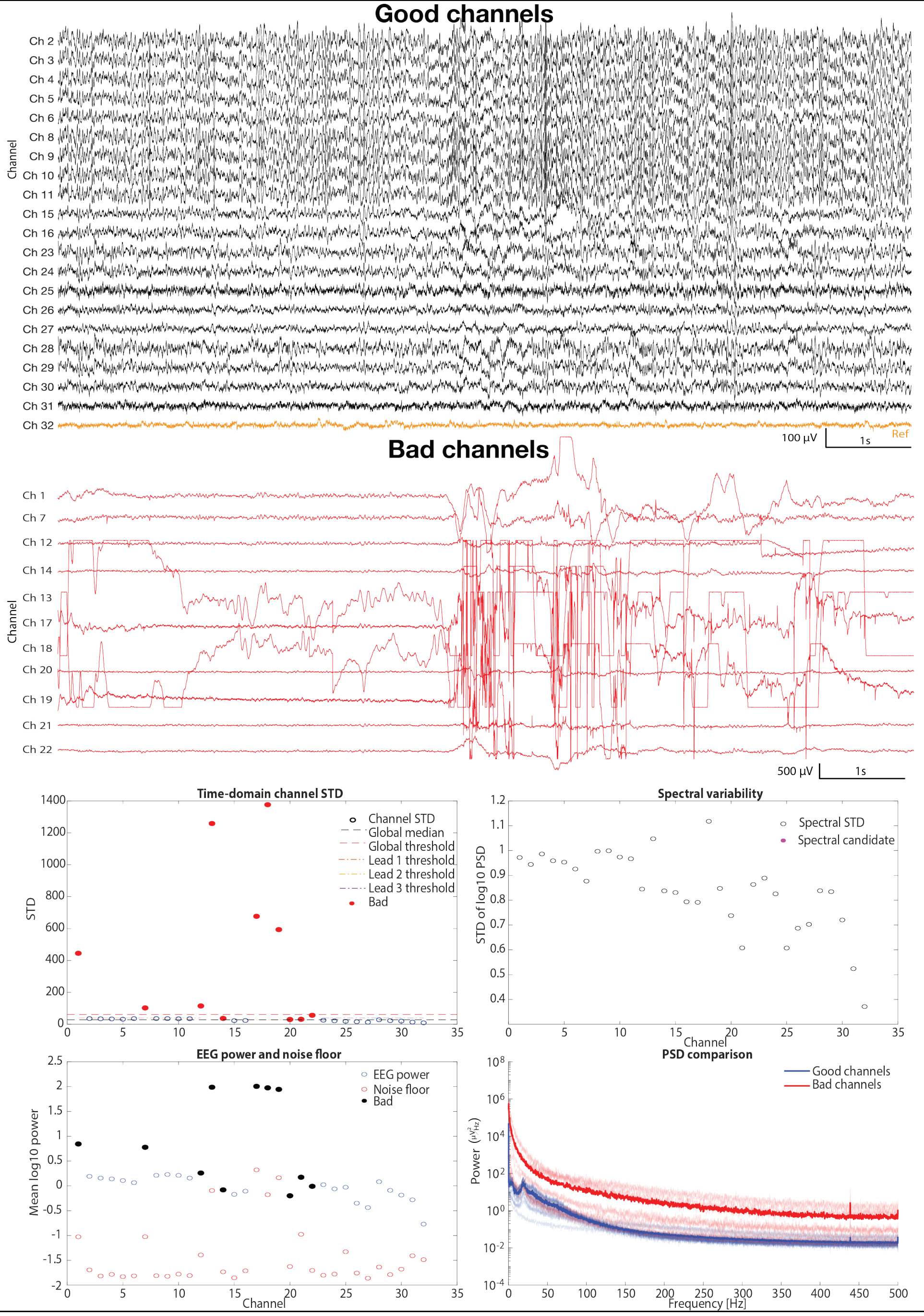

**Figure 13. Detection of bad channels.** Representative good and bad recording channels identified by the automated quality assessment algorithm. Channel classification was based on multiple signal quality metrics, including time-domain signal variability, spectral variability, EEG power, noise floor, power spectral density (PSD), and electrode impedance.

#### Closed-Loop Pipelines

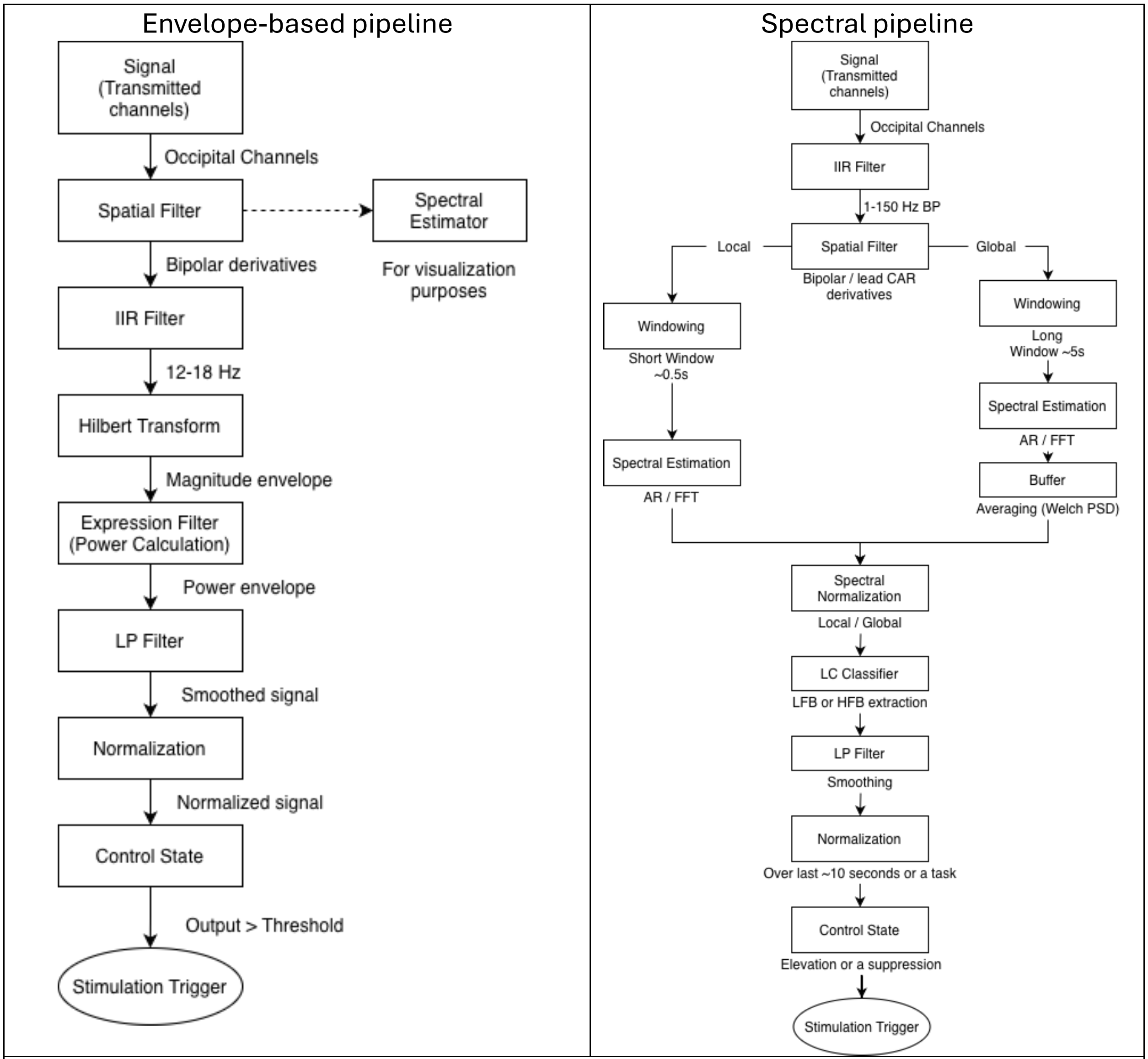

**Figure 14. BCI2000 online signal processing pipelines for closed-loop stimulation**. **Left**: Envelope-based pipeline, in which band-pass filtered neural activity is transformed into an amplitude envelope for real-time detection. This approach offers low latency but reduced specificity. **Right**: Spectral pipeline, in which neural activity is quantified using windowed spectral estimation and normalization. The longer analysis windows increase detection latency but improve specificity and reduce false-positive detections

#### Spike distribution

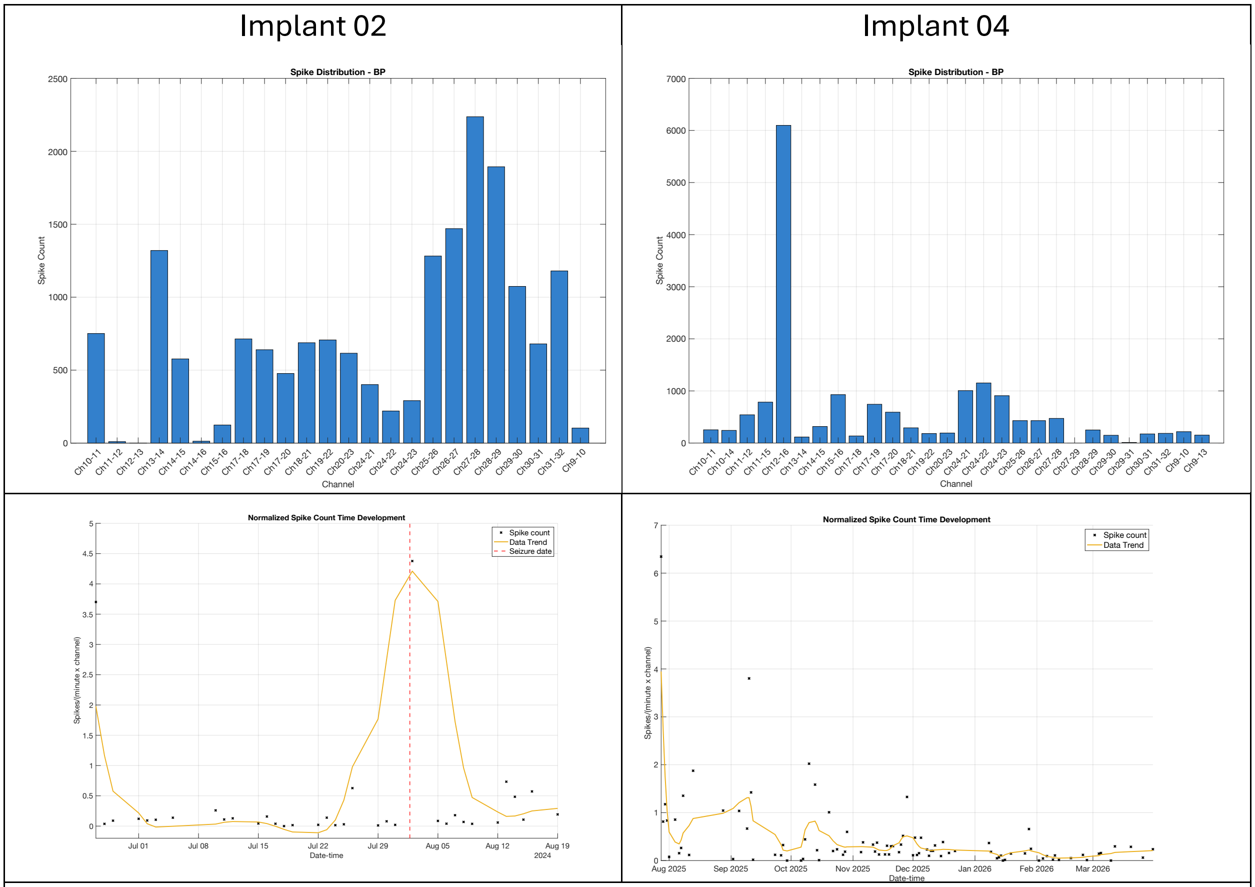

**Figure 15. Longitudinal spike activity in chronic recordings**. **Top**: Distribution of detected spikes across bipolar channel pairs for Implant #2 (left) and Implant #4 (right, epileptic canine). **Bottom:** Longitudinal evolution of the normalized spike count for Implant #2 (left) and Implant #4 (right). The yellow curve represents the smoothed trend in spike activity over time. The dashed vertical line indicates the occurrence of a clinical seizure in Implant #2.

#### Next-generation recording architecture

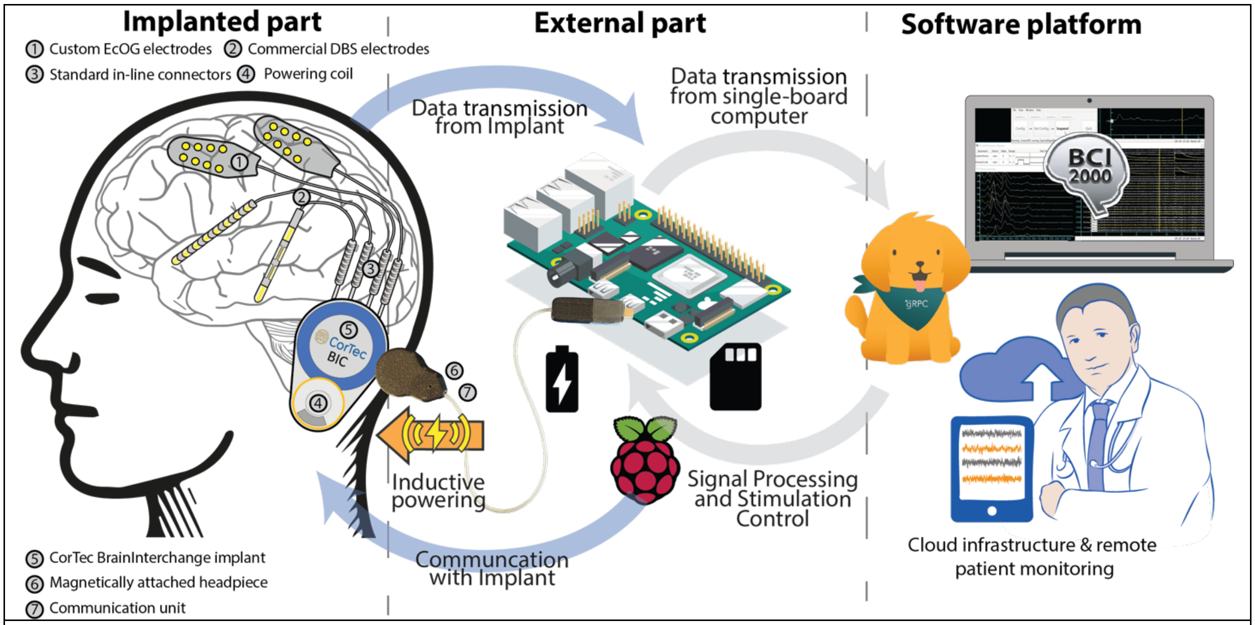

**Figure 16. Next-generation recording architecture.** The proposed BIC ecosystem comprises an implanted neural interface, a miniaturized external communication unit integrated with a portable single-board computer for real-time signal processing and stimulation control, and a software platform supporting BCI2000 integration, cloud connectivity, and remote patient monitoring.
